# Data-Driven Analysis of Age, Sex, and Tissue Effects on Gene Expression Variability in Alzheimer’s Disease

**DOI:** 10.1101/498527

**Authors:** Lavida R.K. Brooks, George I. Mias

## Abstract

Alzheimer’s disease (AD) has been categorized by the Centers for Disease Control and Prevention (CDC) as the 6^th^ leading cause of death in the United States. AD is a significant health-care burden because of its increased occurrence (specifically in the elderly population) and the lack of effective treatments and preventive methods. With an increase in life expectancy, the CDC expects AD cases to rise to 15 million by 2060. Aging has been previously associated with susceptibility to AD, and there are ongoing efforts to effectively differentiate between normal and AD age-related brain degeneration and memory loss. AD targets neuronal function and can cause neuronal loss due to the buildup of amyloid-beta plaques and intracellular neurofibrillary tangles.

Our study aims to identify temporal changes within gene expression profiles of healthy controls and AD subjects. We conducted a meta-analysis using publicly available microarray expression data from AD and healthy cohorts. For our meta-analysis, we selected datasets that reported donor age and gender, and used Affymetrix and Illumina microarray platforms (8 datasets, 2,088 samples). Raw microarray expression data were re-analyzed, and normalized across arrays. We then performed an analysis of variance, using a linear model that incorporated age, tissue type, sex, and disease state as effects, as well as study to account for batch effects, and including binary interaction between factors. Our results identified 3,735 statistically significant (Bonferroni adjusted p<0.05) gene expression differences between AD and healthy controls, which we filtered for biological effect (10% two-tailed quantiles of mean differences between groups) to obtain 352 genes. Interesting pathways identified as enriched comprised of neurodegenerative diseases pathways (including AD), and also mitochondrial translation and dysfunction, synaptic vesicle cycle and GABAergic synapse, and gene ontology terms enrichment in neuronal system, transmission across chemical synapses and mitochondrial translation.

Overall our approach allowed us to effectively combine multiple available microarray datasets and identify gene expression differences between AD and healthy individuals including full age and tissue type considerations. Our findings provide potential gene and pathway associations that can be targeted to improve AD diagnostics and potentially treatment or prevention. (US).

## 1 INTRODUCTION

Aging refers to the physiological changes that occur within the body overtime (Lopez-Otin et al., 2013). These changes are accompanied by deteriorating cell and organ function due to cellular and immune senescence and DNA and protein damage (Lopez-Otin et al., 2013; Van Deursen, 2014; Childs et al., 2015). Aging causes an increased risk for diseases. Age-related diseases are becoming a public health concern due to an overall increase in the older population and the average human life span in developed countries (Rowe et al., 2016; Black et al., 2015). It is predicted that by the year 2050, the number of Americans over 85 years of age will triple from 2015 (Jaul and Barron, 2017; United Nations Department of Economic and Social Affairs, 2015). Larger percentages of the elderly and their increased risk for diseases can affect the economy, social and health care costs (Dallmeyer et al., 2017). For instance, immune system dysfunction and cognitive decline due to aging increases the risk of neurodegenerative diseases such as Alzheimer’s disease (AD) (Jevtic et al., 2017; Mattson and Arumugam, 2018). Previous research explored brain aging and found notable changes in brain size, brain structure and function (Drayer, 1988). Changes in the brain due as we age are also known as hallmarks of brain aging. These hallmarks include: mitchondrial dysfunction, damage to proteins and DNA due to oxidation, neuroinflammation due to immune system dyfunction, reduction in brain volume size and gray and white matter, impaired regulation of neuronal Ca^2+^ (Mattson and Arumugam, 2018; Drayer, 1988). These alterations render the aging brain vulnerable to neurodegenerative diseases such as AD.

AD, the most common form of dementia, is currently the 6^th^ leading cause of death (Taylor et al., 2017) in the United States (US. In 2010, an estimate of 4.7 million people in the US had AD, and the number of AD patients is expected to increase to 13.8 million in 2050 and to 15 million by 2060 (Matthews et al., 2018; Brookmeyer et al., 2018; Hebert et al., 2013). As with other age-related diseases, the risk of AD increases with age. AD is currently characterized by the accumulation of amyloid-beta (Aβ) plaques and neurofibrillary tangles due to tau protein modifications (Masters et al., 2015). These two protein changes are the main pathological changes in AD (Masters et al., 2015). Aβ is formed when the amyloid precursor protein (APP) is cleaved by γ-secretases and β-secretases. Cleavage of APP forms fragments of Aβ which aggregate and deposit on neurons as plaques, which causes neuronal death in conjunction with neurofibrillary tangles (Masters et al., 2015).

While AD’s prevalence is on the rise due to increased life expectancy, there is still no treatment available and diagnosis of AD is challenging. How AD progresses is still not completely understood (De Jager et al., 2018). New technologies are available such as positron-emission tomography (PET) imaging and monitoring levels of Aβ and tau in cerbrospinal fluid (Masters et al., 2015). Co-morbidities that can exist due to aging such as hippocampal sclerosis further complicate AD diagnosis(Toepper, 2017). Furthermore, questions have been raised regarding whether or not AD is simply an accelerated form of aging due to them both being associated with changes in cognition (Toepper, 2017). However, studies have identified clear neurocognitive differences in cognition, brain size and function in AD compared to healthy aged subjects. For example, AD patients have more grey matter loss compared to white matter, impaired verbal and semantic abilities and more intense memory dysfunction compared to healthy seniors (Toepper, 2017).

Pathological changes within the brain are observed prior to clinical diagnosis of AD. In most cases AD cannot be confirmed until postmortem examination of the brain. Researchers are investigating novel biomarkers to detect for earlier diagnosis before diseased individuals become functionally impaired. Metaanalysis of microarray datasets is becoming more popular for it provides stronger power to studies due to larger sample sizes, obtained through statistically combining multiple datasets. Microarray data are also available in large quantities on public online data repositories. In the case of AD, Winkler et al., performed a meta-analysis that compared neurons within the hippocampus of AD patients and healthy controls. They identified that processes such as apoptosis, and protein synthesis, were affected by AD and were regulated by androgen and estrogen receptors(Winkler and Fox, 2013). Researchers have also explored differences in gene expression in Parkinson’s and AD subjects via a meta-analysis approach (Wang et al., 2017), and identified functionally enriched genes and pathways that showed overlap between the two diseases (Wang et al., 2017). Most recently, Moradifard et al. identified differentially expressed microRNAs and genes when comparing AD to healthy controls via a meta-analysis approach. They also identified two key microRNAs that act as regulators in the AD gene network(Moradifard et al., 2018).

In our investigation, our goal was to identify age, sex, and tissue effects on gene expression variability in AD by comparing age-matched healthy controls to AD subjects via a meta-analysis approach. In this data-driven approach, we explored global gene expression changes in 2,088 total samples (771 healthy, 868 AD, and 449 possible AD, curated from 8 studies) from 26 different tissues, to identify genes and pathways of interest in AD that can be affected by factors such as age,sex and tissue. Our findings provide potential gene and pathway associations that can be targeted to improve AD diagnostics and potentially treatment or prevention.

## 2 METHODS

We conducted a meta-analysis using 8 publicly available microarray expression datasets (Table 1) from varying tissues and microarray platforms on AD. We developed a thorough computational pipeline (Figure 1A) that involved curating and downloading raw microarray expression data, pre-processing the raw expression data and conducting a linear model analysis of the gene expression profiles. Statistically different genes based on disease state were identified following analysis of variance (ANOVA) on the linear model which compared gene expression changes due to disease state, sex, age and tissue. These genes were further analyzed using a Tukey Honest Significant Difference (TukeyHSD) test to determine their biological significance (Tukey, 1949). In addition to the p-values, we also obtained the mean differrences between binary comparisons of groups (also generated by the TukeyHSD), as a measure of biological effect size. We examined the TukeyHSD results by filtering by each factor, and identified up and down regulated genes. Using these genes we used R packages ReactomePA (Yu and He, 2016) and clusterProfiler (Yu et al., 2012) to conduct gene enrichment and pathway analyses of the differentially expressed genes. We used BINGO in Cytoscape v.3.7.0 for gene ontology (GO) analysis on each gene set for each factor (Maere et al., 2005; Shannon et al., 2003).

**Table 1.** Curated microarray datasets and the study description.

| Database | Accession Number | Controls | AD | Possible AD | Platform | Citation |
| --- | --- | --- | --- | --- | --- | --- |
| GEO | GSE84422 | 242 | 362 | 449 | Affymetrix Human Genome U133A, B and Plus 2.0 | (Wang et al., 2016) |
| GEO | GSE28146 | 8 | 22 | - | Affymetrix Human Genome Plus 2.0 | (Blalock et al., 2011) |
| GEO | GSE48350 | 173 | 80 | - | Affymetrix Human Genome Plus 2.0 | (Berchtold et al., 2008) |
| GEO | GSE5281 | 74 | 85 | - | Affymetrix Human Genome Plus 2.0 | (Liang et al., 2007) |
| GEO | GSE63060 | 104 | 142 | - | Illumina HumanHT-12 V3.0 expression beadchip | (Sood et al., 2015) |
| GEO | GSE63061 | 134 | 139 | - | Illumina HumanHT-12 V4.0 expression beadchip | (Sood et al., 2015) |
| GEO | GSE29378 | 32 | 31 | - | Illumina HumanHT-12 V3.0 expression beadchip | (Miller et al., 2013) |
| Array Express | E-MEXP-2280 | 5 | 7 | - | Affymetrix Human Genome Plus 2.0 | (Bronner et al., 2009) |

**Figure 1.**
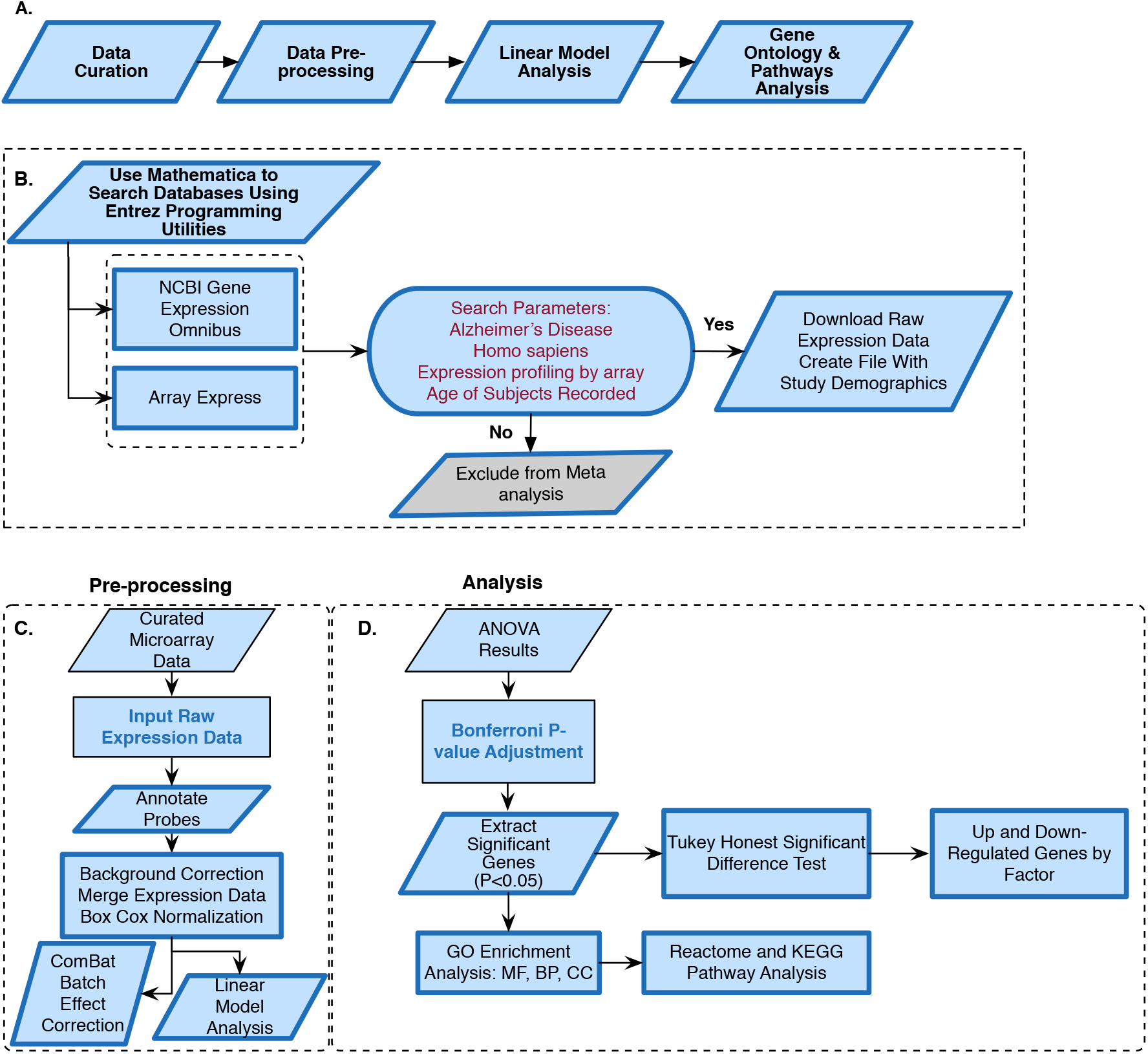
Alzheimer’s disease meta-analysis framework. **(A)** Simplified workflow used for the metaanalysis, **(B)** Pipeline for curating microarray data, **(C)** Pipeline for pre-processing the microarray data, **(D)** Methods used for meta-analysis of raw expression microarray data.

### Microarray Data Curation

We curated microarray expression data from two data repositories: National Center for Biotechnology Information (NCBI) Gene Expression Omnibus (GEO) (Edgar et al., 2002) and Array Express (Brazma et al., 2003) (Figure 1B). We searched these repositories by using entrez programming utilities in Mathematica (Mias, 2018b). In this search, we used the following keywords: *Homo sapiens*, Alzheimer’s Disease and expression profiling by array (Figure 1B). This search resulted in 105 datasets from GEO and 8 from Array Express. We further filtered the search results by excluding data from cell lines, selecting for expression data from Illumina and Affymetrix microarray platforms, and focusing on datasets that provided the ages of their samples (Figure 1B). After filtering through the databases, we found 7 datasets from GEO (GSE84422, GSE28146, GSE48350, GSE5281,GSE63060,GSE63061,GSE29378) and 1 dataset from Array Express (E-MEXP-2280) to conduct our meta-analysis of expression profiling to assess differences in gene expression due to disease state, sex, age and tissue (Table 1). Additionally, we downloaded the raw expression data from each dataset, and created a demographics file per study which included characteristics about the samples (Table 2). To ensure uniform annotation of the subjects we made some changes to subject information provided from the databases. For GSE28146, we grouped the sub-types of AD: incipient, moderate and severe as AD because we did not have classification information for our other AD samples. We changed all the GSE29378 tissue types to hippocampus, relabeled the “probable AD” disease state to “possible AD” in GSE84422, only used AD and control subjects from the E-MEXP-2280 and GSM238944 with an age of >90 (not a definite age) was removed from GSE5281. We should note also that the 1,053 samples from the GSE84422 dataset included different tissues from the same subjects.

**Table 2.** Patient characteristics for curated datasets.

| Accession Number | Sex (M/F) | Age Range |
| --- | --- | --- |
| GSE84422 | 302M/166F | 60-103 |
| GSE28146 | 12M/18F | 65-101 |
| GSE48350 | 124M/129F | 20-99 |
| GSE5281 | 102M/56F | 63-102 |
| GSE63060 | 88M/158F | 52-88 |
| GSE63061 | 107M/166F | 59-95 |
| GSE29378 | 38M/25F | 61-90 |
| E-MEXP-2280 | 7M/5F | 68-82 |

### Pre-processing and Data Normalization

We downloaded the raw expression data from the database repositories in Mathematica (Wolfram Research, Inc., 2017) and pre-processed each file in R (R Core Team, 2018) using the appropriate R packages based on the microarray platform. The affy package was used to pre-process all the .CEL data files from affymetrix, and limma for Illumina summary data files. We performed background correction, normalization and annotated and summarized all probes (Figure 1C). We merged all 8 datasets into one large matrix file via common gene names. After merging the datasets, we performed a BoxCox power transformation (Sakia, 1992) and data standardization in Mathematica (Figure 1C) (see ST2 of online supplemental data).

### Correcting for Batch Effects

Merging expression data from different studies, array platforms and tissues can introduce confounding factors and manipulate interpretation of results. To address this, and assess whether batch effects were evident and could be accounted for, we used the ComBat (Nygaard et al., 2016) (Johnson et al., 2007) algorithm to adjust data for known batch effects. In this study, the batch effect was the study (i.e. different experiments/groups), and we also noted that there was a one-to-one correspondence between study and platform. Using data from prior to and post ComBat corrections, we used principal component analysis (PCA) plots to visualize the variability in the data and the effectiveness of possible batch effect removal (Irizarry and Love, 2015).

### Linear Model Analysis and Analysis of Variance

We modeled the merged expression data (see model breakdown below) prior to running analysis of variance (ANOVA) to analyze differences among the different study factors (Figure 1D) (Pavlidis, 2003). We defined age group, sex, disease state, study and tissue as factors.

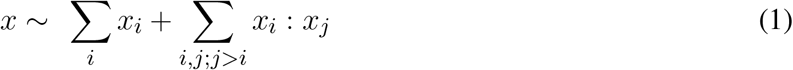

where *x*_*i*_ ∈ {age group, sex, tissue, disease status} and the factors have the following levels

- disease status = {control, possible AD, AD}
- sex = {male, female}
- age group = {under 60, 60-65, 65-70, 70-75, 75-80, 80-85, 85-90, 90-95, over 95}
- tissue = {amygdala, anterior cingulate, blood, caudate nucleus, dorsolateral prefrontal cortex, entorhinal cortex, frontal pole, hippocampus, inferior frontal gyrus, inferior temporal gyrus, medial temporal lobe, middle temporal gyrus, nucleus accumbens, occipital visual cortex, parahippocampal gyrus, posterior cingulate cortex, precentral gyrus, prefrontal cortex, primary visual cortex, putamen, superior frontal gyrus, superior parietal lobule, superior temporal gyrus, temporal pole}
- study = {GSE84422, GSE28146, GSE48350, GSE5281, GSE63060, GSE63061, GSE29378, E-MEXP-2280}

The p-values following the ANOVA were adjusted using Bonferroni correction for multiple hypothesis testing (Pavlidis, 2003). Genes with p-values <0.05 were considered statistically significant. We found significantly different disease genes by filtering on the disease status for p-values <0.05. Additionally, we used the clusterprofiler package in R for Kyoto Encyclopedia of Genes and Genomes (KEGG) enrichment analysis on these genes (Yu et al., 2012; Kanehisa and Goto, 2000). This package adjusts p-values using the Benjamini Hochberg method for False Discovery Rate (FDR) control. Pathways with adjusted p-value <0.05 were considered significantly enriched (Yu et al., 2012).

### Identifying Up and Down Regulated Genes by Factor

To identify which of the 3,735 genes that show biologically significant differences, we conducted a Tukey Honest Significant Difference (TukeyHSD) test to determine statistically significant up and down-regulated genes using the difference in the means of pairwise comparisons between the levels within each factor (Tukey, 1949; Mias, 2018a). We carried out TukeyHSD testing on the statistically significantly different disease genes we obtained from the ANOVA. To account for multiple hypothesis testing in the TukeyHSD results, we used <0.00013 (0.05/number of genes ran through TukeyHSD) as a Bonferroni adjusted cutoff for significance.

We selected the TukeyHSD results from the disease status factor, and focused on the “Control-AD” pairwise comparison to assess statistically significant gene expression differences. To assess biological effect, and select an appropriate fold-change-like cutoff (as our results had already been transformed using a Box-Cox transformation), we calculated the quantiles based on the TukeyHSD difference of mean difference values (Supplemental Table 1). We used a two-tailed 10% and 90% quantile to identify significantly up and down regulated genes (Supplemental Table 1).

The differentially expressed genes (DEG) by disease status factor were subsequently used to determine whether or not there was a sex, age or tissue effect on them. For sex, we used the DEG to filter the TukeyHSD results for sex factor differences, identified statistically significant sex-relevant genes based on p-value cutoff, and again computed the 10% and 90% quantile based on the difference of means between male and female groups. We repeated the above steps for age group, but focused only on the binary comparisons where all age groups were compared to the <60 age group, which was used as a baseline (i.e. computed the mean gene expression differences per group comparison, *i*-< 60, where *i* stands for any age group). This was carried out to enable us to compare the progression with age, relative to a common reference across all age groups. As for tissue, we carried out the same steps as above to determined DEG based on comparisons both a hippocampus-based baseline, as well a blood-based baseline.

Following the identification of the DEG by disease status and sex, we visualized the raw expression data for these genes in heatmaps. In addition to this, we generated heatmaps using the difference of means values for the significantly different age group and tissue genes.

### Gene Ontology and Reactome Pathway Analysis

For the disease and sex DEG sets, we used the R package reactomePA to find enriched pathways(Yu and He, 2016). We also built networks to determine if genes overlapped across pathways. Additionally, we used BINGO in Cytoscape for GO analysis to determine the biological processes the genes were enriched in (Maere et al., 2005). Results were considered significant based on Benjamini-Hochberg adjusted p-value <0.05.

## 3 RESULTS

With our data selection criteria outlined in Figure 1B we identified 8 datasets from GEO and Array Express to conduct our meta-analysis to assess differences in gene expression due to disease state, sex, age and tissue (Table 1). We merged the processed expression data by common gene names, which gave us a total of 2,088 samples and 16,257 genes. The 2,088 samples consisted of 771 healthy controls, 868 AD subjects, 449 subjects reported as possibly having AD, 1308 females and 780 males.

### ComBat Batch Effect Correction

Combining data from different platforms, tissues and different laboratories introduces batch effects. Batch effects are sources of non-biological variations that can affect conclusions. We used ComBat in R which works by adjusting the data based on a known batch effect. For our analysis we classified the study variable as our batch (and also the study and type of platform are directly related). We used PCA to visualize variation in the merged expression data before and after ComBat. In Figure 2 before correcting for batch effects, the datasets separate into 4 main clusters with a variance of 54.3% in PC1 and 13% in PC2. Following ComBat, those main clusters appear to be removed, with an overall reduction in variation for both principal components. We also looked at how the data separated by factor. In Figure 2B, there are two clear groups and this separation is accounted for when we look at the separation in the data by tissue (Figure 3). In Figure 3, before correction the 4 groups observed in Figure 2 are still evident. Following ComBat, the tissues: amygdala and nucleus accumbens cluster together in one group while all other tissues are in another. Visualizing and understanding the variation within the expression data following the merge confirmed the need to include the study as a factor in the linear model analysis.

**Figure 2.**
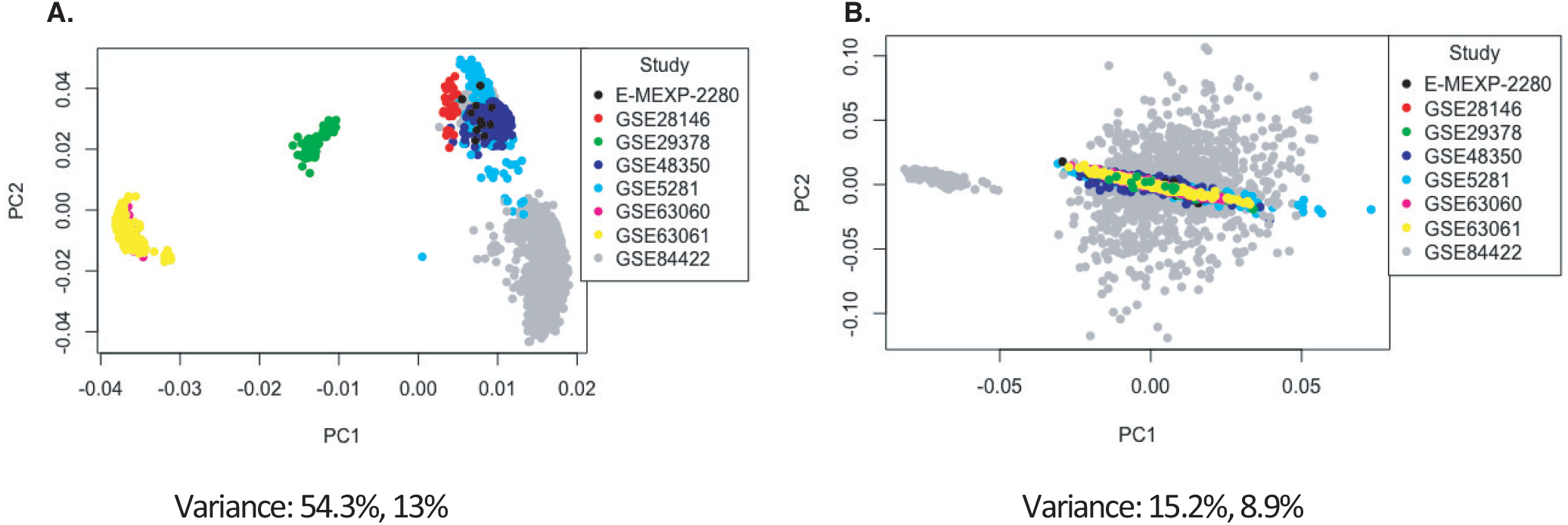
Principal component analysis of the study factor before and after batch correction with ComBat.

**Figure 3.**
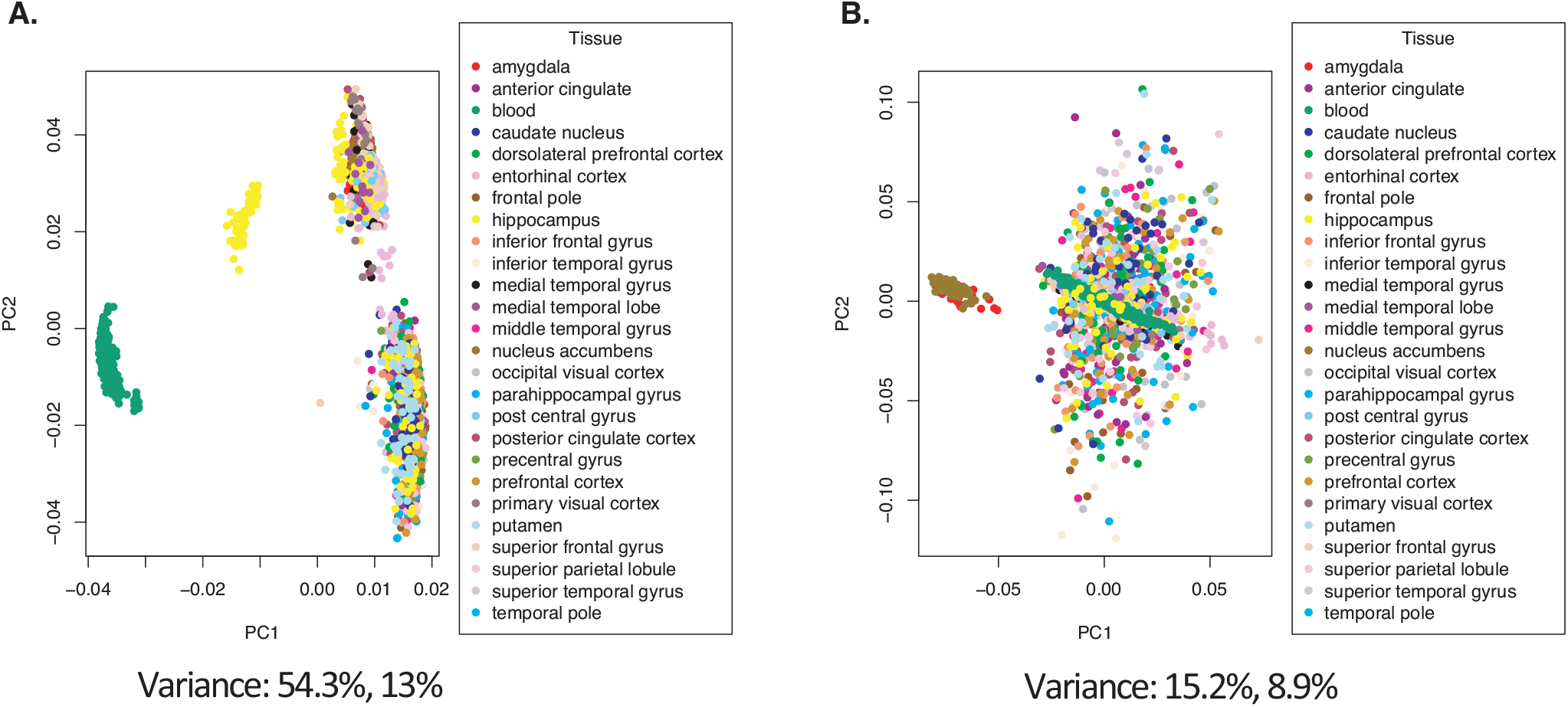
Principal component analysis of the tissue factor before and after batch correction with ComBat.

### Analysis of Variance on Gene Expression By Disease State

Using ANOVA we assessed the variance in gene expression across the different factors in our linear model by including the following factors and their pairwise interactions: age group, study, tissue, sex and disease state (Pavlidis, 2003). Statistically significantly gene expression differences were determined using a Bonferroni((Bland and Altman, 1995) adjusted p-value was (<0.05) (Pavlidis, 2003; Mias, 2018a). With our focus on differences by disease status, we filtered genes based on the ANOVA adjusted p-values for the disease factor. Selecting for significance by disease status we found 3,735 genes (see ST4 of online supplemental data). We conducted GO and pathway analysis on these genes. The KEGG pathway analysis results are displayed in Table 3 (see ST5 of online supplemental data for full table). The analysis showed that the genes are involved in Reactome pathways such as the Mitochondrial Translation Initiation (55 gene hits), Signaling by the B Cell Receptor (61 gene hits), Activation of NF-kappaβin B cells (40 gene hits), Transmission across Chemical Synapses (83 gene hits) and Neuronal System (119 gene hits) (see ST6 of online supplemental data). The KEGG pathways that were enriched for this gene set included neurodegenerative disease pathways such as Alzheimer’s (31 gene hits), Huntington’s (76 gene hits) and Parkinson’s (53 gene hits) (Table 3) Pathways. There were also some synaptic pathways: Synaptic vesicle cycle (30 gene hits), Dopaminergic synapse (48 gene hits) and GABAergic synapse (34 gene hits) (Table 3). In addition to synapses and neurodegeneration, the long term potentiation (23 gene hits) pathway was associated with these genes (see ST5 for full KEGG pathway analysis results). To further explore the enriched genes in the KEGG AD pathway, we used the TukeyHSD results to determine whether genes were up- or down- regulated (see ST7 of online supplemental data). To further assess the 73 gene hits identified in the enriched AD pathway we computed their mean differences between the AD and control, and used MathIOmica (Mias et al., 2016) tools to highlight them in the AD pathway (Figure 4) (Mias, 2018b; Kanehisa and Goto, 2000; Kanehisa et al., 2016, 2017) (see ST7 on online supplement data for full table with difference of means). For instance, APOE gene in the AD pathway is down-regulated in healthy controls compared to AD subjects (Figure 4).

**Table 3.**
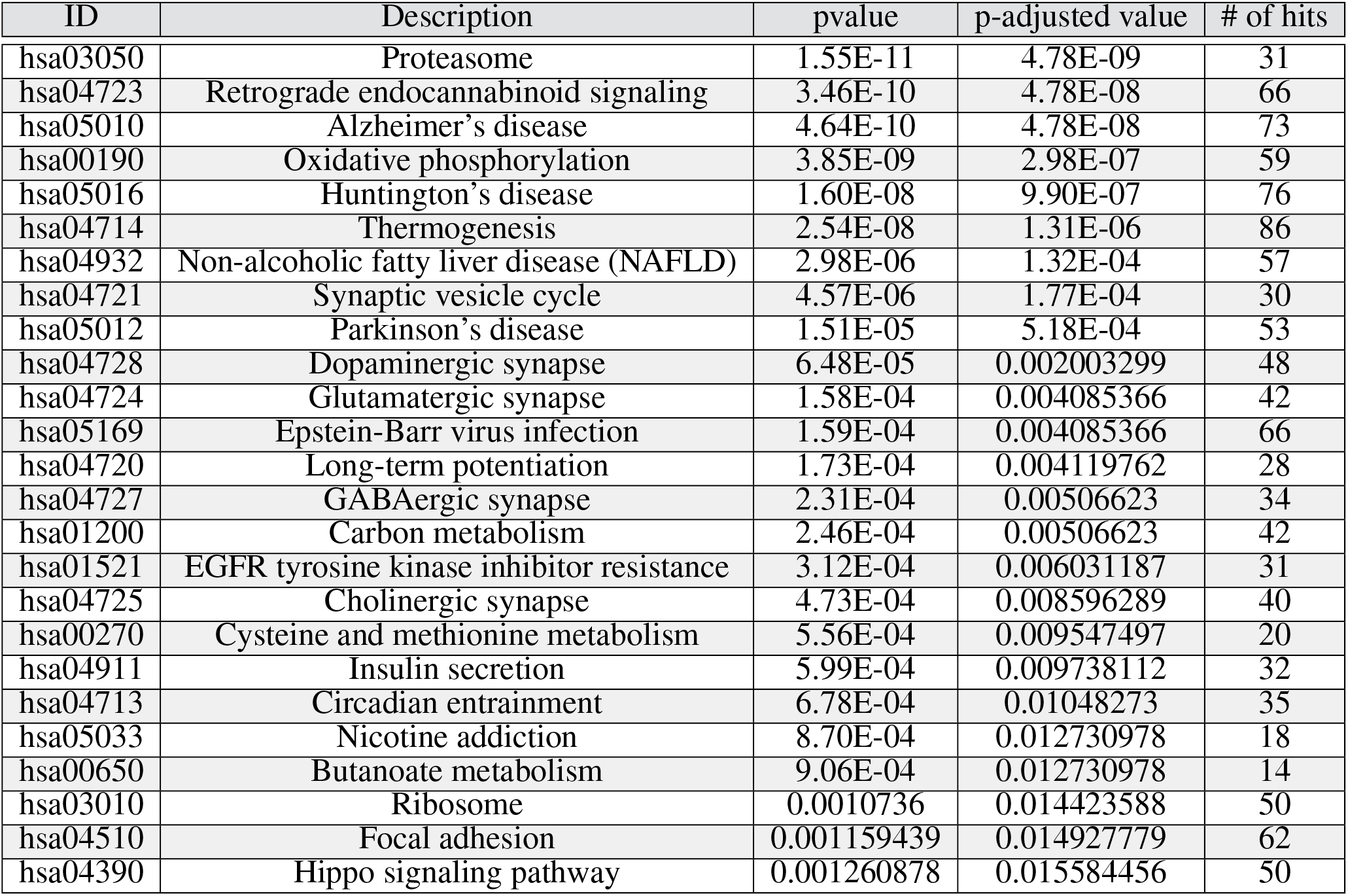
Top 25 KEGG Pathways using differentially expressed genes.

| ID | Description | pvalue | p-adjusted value | # of hits |
| --- | --- | --- | --- | --- |
| hsa03050 | Proteasome | 1.55E-11 | 4.78E-09 | 31 |
| hsa04723 | Retrograde endocannabinoid signaling | 3.46E-10 | 4.78E-08 | 66 |
| hsa05010 | Alzheimer's disease | 4.64E-10 | 4.78E-08 | 73 |
| hsa00190 | Oxidative phosphorylation | 3.85E-09 | 2.98E-07 | 59 |
| hsa05016 | Huntington's disease | 1.60E-08 | 9.90E-07 | 76 |
| hsa04714 | Thermogenesis | 2.54E-08 | 1.31E-06 | 86 |
| hsa04932 | Non-alcoholic fatty liver disease (NAFLD) | 2.98E-06 | 1.32E-04 | 57 |
| hsa04721 | Synaptic vesicle cycle | 4.57E-06 | 1.77E-04 | 30 |
| hsa05012 | Parkinson's disease | 1.51E-05 | 5.18E-04 | 53 |
| hsa04728 | Dopaminergic synapse | 6.48E-05 | 0.002003299 | 48 |
| hsa04724 | Glutamatergic synapse | 1.58E-04 | 0.004085366 | 42 |
| hsa05169 | Epstein-Barr virus infection | 1.59E-04 | 0.004085366 | 66 |
| hsa04720 | Long-term potentiation | 1.73E-04 | 0.004119762 | 28 |
| hsa04727 | GABAergic synapse | 2.31E-04 | 0.00506623 | 34 |
| hsa01200 | Carbon metabolism | 2.46E-04 | 0.00506623 | 42 |
| hsa01521 | EGFR tyrosine kinase inhibitor resistance | 3.12E-04 | 0.006031187 | 31 |
| hsa04725 | Cholinergic synapse | 4.73E-04 | 0.008596289 | 40 |
| hsa00270 | Cysteine and methionine metabolism | 5.56E-04 | 0.009547497 | 20 |
| hsa04911 | Insulin secretion | 5.99E-04 | 0.009738112 | 32 |
| hsa04713 | Circadian entrainment | 6.78E-04 | 0.01048273 | 35 |
| hsa05033 | Nicotine addiction | 8.70E-04 | 0.012730978 | 18 |
| hsa00650 | Butanoate metabolism | 9.06E-04 | 0.012730978 | 14 |
| hsa03010 | Ribosome | 0.0010736 | 0.014423588 | 50 |
| hsa04510 | Focal adhesion | 0.001159439 | 0.014927779 | 62 |
| hsa04390 | Hippo signaling pathway | 0.001260878 | 0.015584456 | 50 |

**Figure 4.**
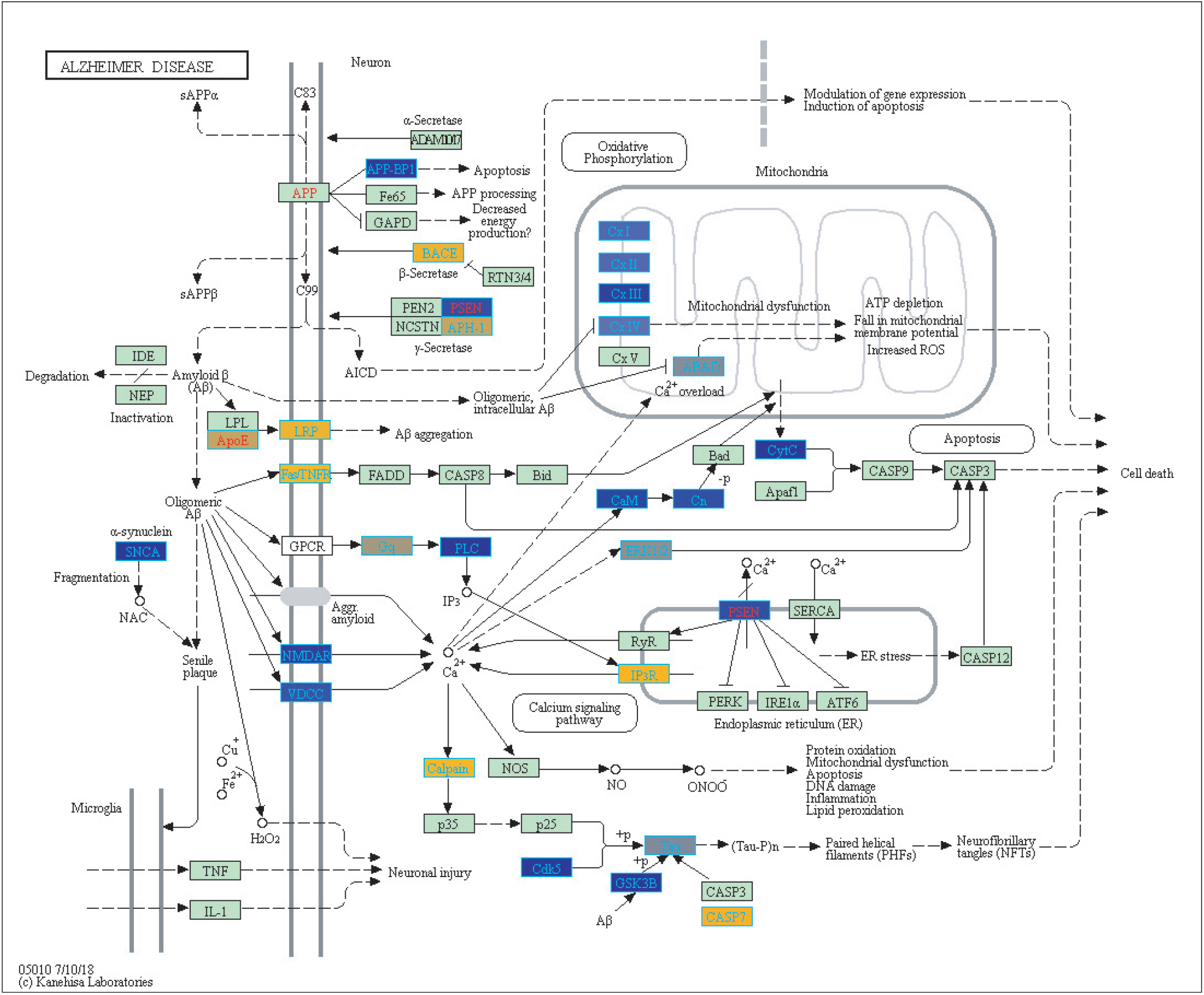
Enriched genes from the ANOVA significant disease gene list in KEGG Alzheimer’s disease pathway (hsa05010) [Kanehisa and Goto (2000); Kanehisa et al. (2016, 2017)]. The yellow shading represents up-regulated and the blue shading represents down-regulated in AD samples.

### Up and Down- Regulated Gene Expression in AD and Sex Specific Differences

We conducted a post-hoc analysis (TukeyHSD) on the 3,735 statistically significant disease genes to identify level differences and explore up- and down- regulation of genes. We were particularly interested in the control compared to AD gene expression differences, and how these could be further sub-categorized to explore effects by sex, age and tissue. We used a Bonferroni adjusted p-value cut off for significance (<0.000013) and the 10% two-tailed quantile to determine significantly up and down regulated genes (Supplemental Table 1). In the Control-AD TukeyHSD comparisons, genes were classified as up- regulated (352 DEG) and down- regulated (176 DEG) in AD (or correspondingly up or down- regulated in healthy) if their mean differences were ≤ −0.0945 and ≥ 0.1196 respectively (Supplemental Table 1, see also ST8 of online supplemental data). The top 25 up- and down- regulated genes sorted by the TukeyHSD adjusted p-values are outlined in Table 4. After performing gene enrichment and pathway analysis with ReactomePA (Yu and He, 2016) on the 352 genes we built pathway-gene networks for the significant Reactome pathways (Benjamini-Hochberg adjusted p-value < 0.05) (see ST13 and ST14 of online supplemental data). Some of the top 10 enriched Reactome pathways from DEG down regulated in AD include: Mitochondrial translation elongation, Mitochondrial translation, Transmission across chemical synapses, neuronal system (Figure 5). The network in Figure 5 illustrates that some genes overlap across pathways - the difference of means from the TukeyHSD results of these genes are indicated by the color scale. The up-regulated genes in AD were enriched in pathways such as Extracellular matrix (ECM) organization and ECM proteoglycans, Non-integrin membrane-ECM interactions and potassium channels. Additionally, we used BINGO for GO analysis on the 352 disease DEG to determine the biological processes they are involved in (Supplemental Figure 6). Some examples of significant terms: Cell signaling development, nervous system development, neuron differentiation, cell proliferation, response to chemical stimulus, cell communication and brain and nervous system development (Supplemental Figure 6).

**Table 4.** Top 25 up- and down- regulated genes in Alzheimer’s disease compared to healthy controls.

| Up-Regulated |  | Down-Regulated |  |
| --- | --- | --- | --- |
| Gene | Difference of Means | Gene | Difference of Means |
| ITPKB | 0.1709575 | RPA3 | -0.1781622 |
| ARHGEF40 | 0.1574220 | NME1 | -0.1755078 |
| CXCR4 | 0.1907433 | LSM3 | -0.1527917 |
| PRELP | 0.1319160 | MRPL3 | -0.1577078 |
| SLC7A2 | 0.1568425 | PTRH2 | -0.1205413 |
| AHNAK | 0.1304494 | RGS7 | -0.1778522 |
| NOTCH1 | 0.1014441 | GLRX | -0.1622333 |
| GFAP | 0.1198343 | RPH3A | -0.2168597 |
| HVCN1 | 0.1151989 | BEX4 | -0.1416335 |
| LDLRAD3 | 0.1627433 | COX7B | -0.1726039 |
| KANK1 | 0.0992824 | NRN1 | -0.1634702 |
| HIPK2 | 0.1255059 | PPEF1 | -0.1430548 |
| SLC6A12 | 0.1485253 | PCSK1 | -0.3127961 |
| KLF4 | 0.1870071 | ENY2 | -0.1496523 |
| ABCA1 | 0.1386346 | CD200 | -0.1537059 |
| DDR2 | 0.1069751 | NRXN3 | -0.1203814 |
| KLF2 | 0.1070143 | GTF2B | -0.1508171 |
| GNG12 | 0.1318200 | MRPS18C | -0.1535766 |
| POU3F2 | 0.1022426 | NCALD | -0.1858802 |
| AEBP1 | 0.1498719 | C11orf1 | -0.1448555 |
| IQCA1 | 0.1134073 | DCTN6 | -0.1222108 |
| ERBIN | 0.1309312 | SEM1 | -0.1765024 |
| LOC202181 | 0.1184466 | APOO | -0.1384320 |
| LPP | 0.1072798 | CCNH | -0.1394853 |
| NOTCH2 | 0.1213843 | RAD51C | -0.1280948 |

**Figure 5.**
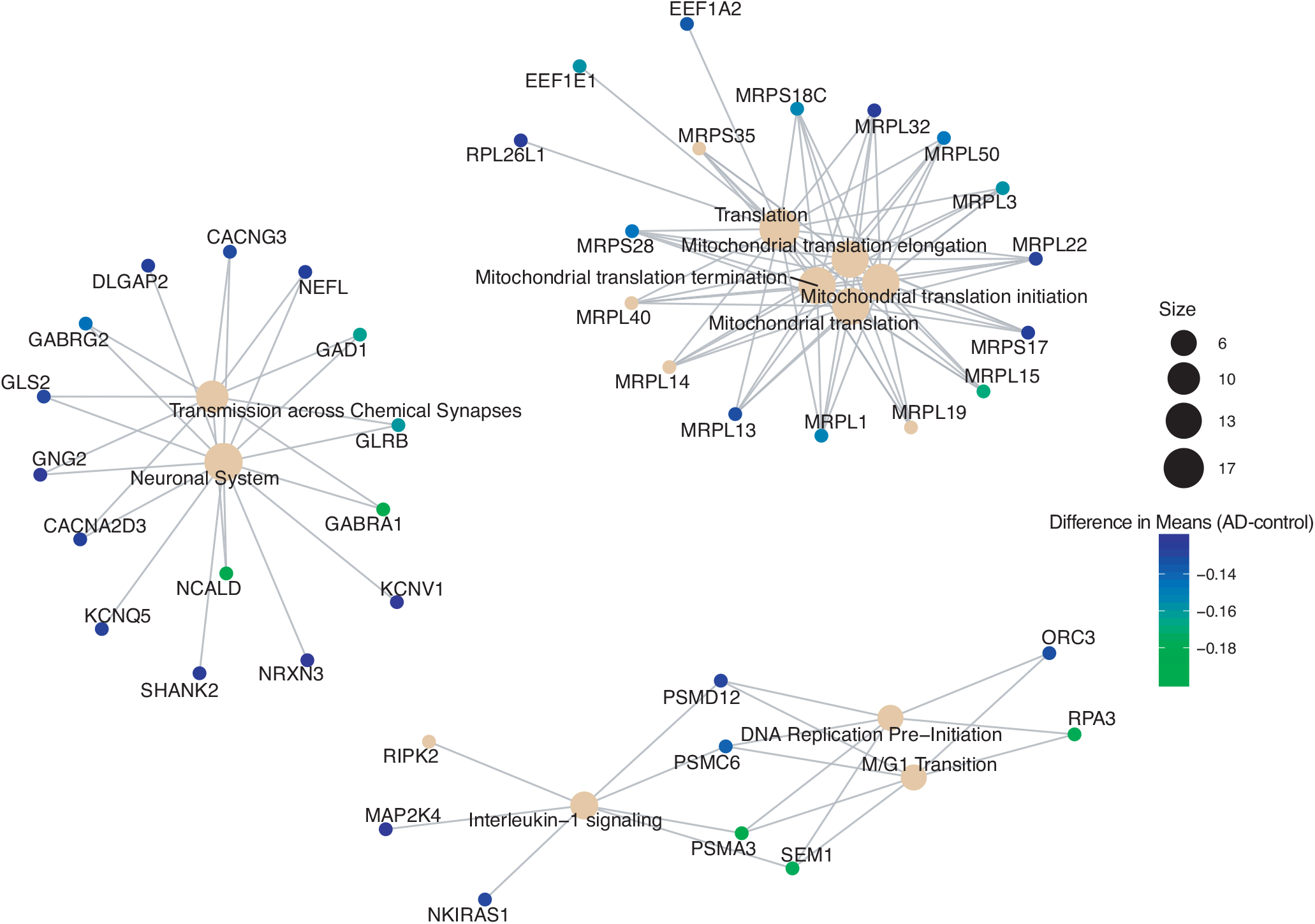
Pathway-gene network of top 10 enriched Reactome pathways from down-regulated genes in Alzheimer’s disease patients.

**Figure 6.**
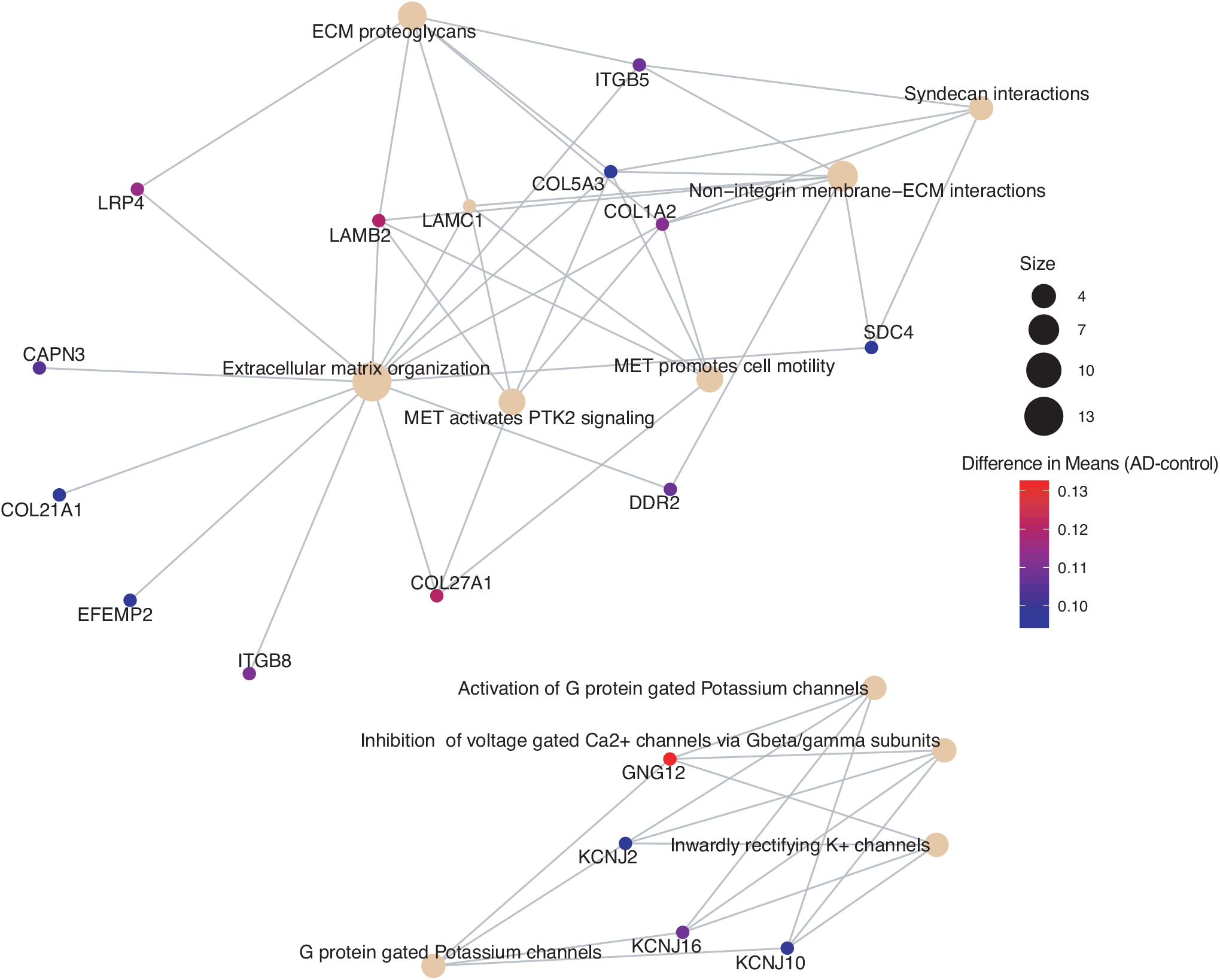
Pathway-gene network of top 10 enriched Reactome pathways from up-regulated genes in Alzheimer’s disease patients.

Of the 352 DEG in the above disease analysis, 46 genes differentially expressed by sex: 23 down- and 23 up- regulated in males compared to females (Supplementary Table 2) based on mean differences (≤ −0.0864 and ≥ 0.2502 respectively (Supplementary Table 1). We used ReactomePA to build a network of enriched genes and pathways with sex differences (Figure 7) (Yu and He, 2016). We found 6 pathways that were enriched with the up-regulated gene list in males: Neuronal System, Transmission across chemical synapses, neurotransmitter receptors and post-synaptic signal transmission, and GABA A receptor activation (Figure 7 - see also ST9 of online supplemental data).

**Figure 7.**
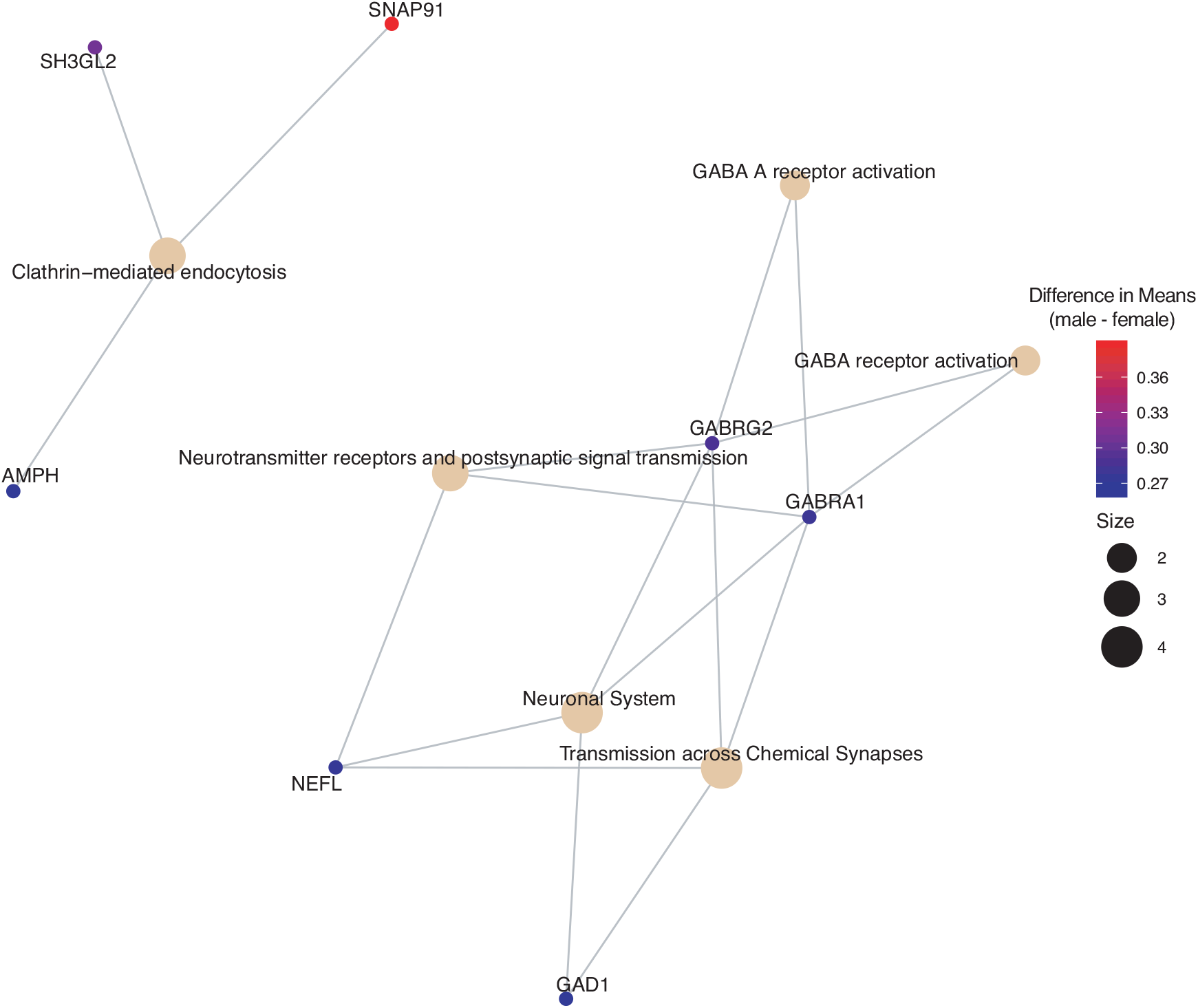
Pathway-gene network of enriched Reactome pathways from disease genes list that are up- regulated in males

### Aging and Tissue Differences in AD Gene Expression

To determine if age or tissue had an effect on the differentially expressed genes by disease status, we filtered the 352 DEG in disease results discussed above for age group and tissue comparisons. For age effects, we used our TukeyHSD results that compared age groups to <60 (served as the baseline). This allowed us to explore if genes associated with AD change with age by using a common reference group. We used the 352 DEG genes from disease status TukeyHSD results to find sizable age effects in this gene set by selecting for statistical significance and using the two-tailed 10% quantile filter (≤ −1.0477827 and ≥ 0.330869) to find significantly different genes per age-group pair comparison (Supplemental Table 1). We found 396 significant comparisons of age differences in 141 genes (see ST10 of online supplemental data). The 141 genes were plotted across all age comparisons where < 60 was the baseline to visualize expression changes and how the genes clustered(Figure 8), indicative of distinct differences in expression profiles due to aging. There is a cluster of genes down-regulated in older age groups, specifically ages 65-80 compared to those < 60. There also appears to be an overall trend of genes associated with disease being up-regulated compared to < 60.

**Figure 8.**
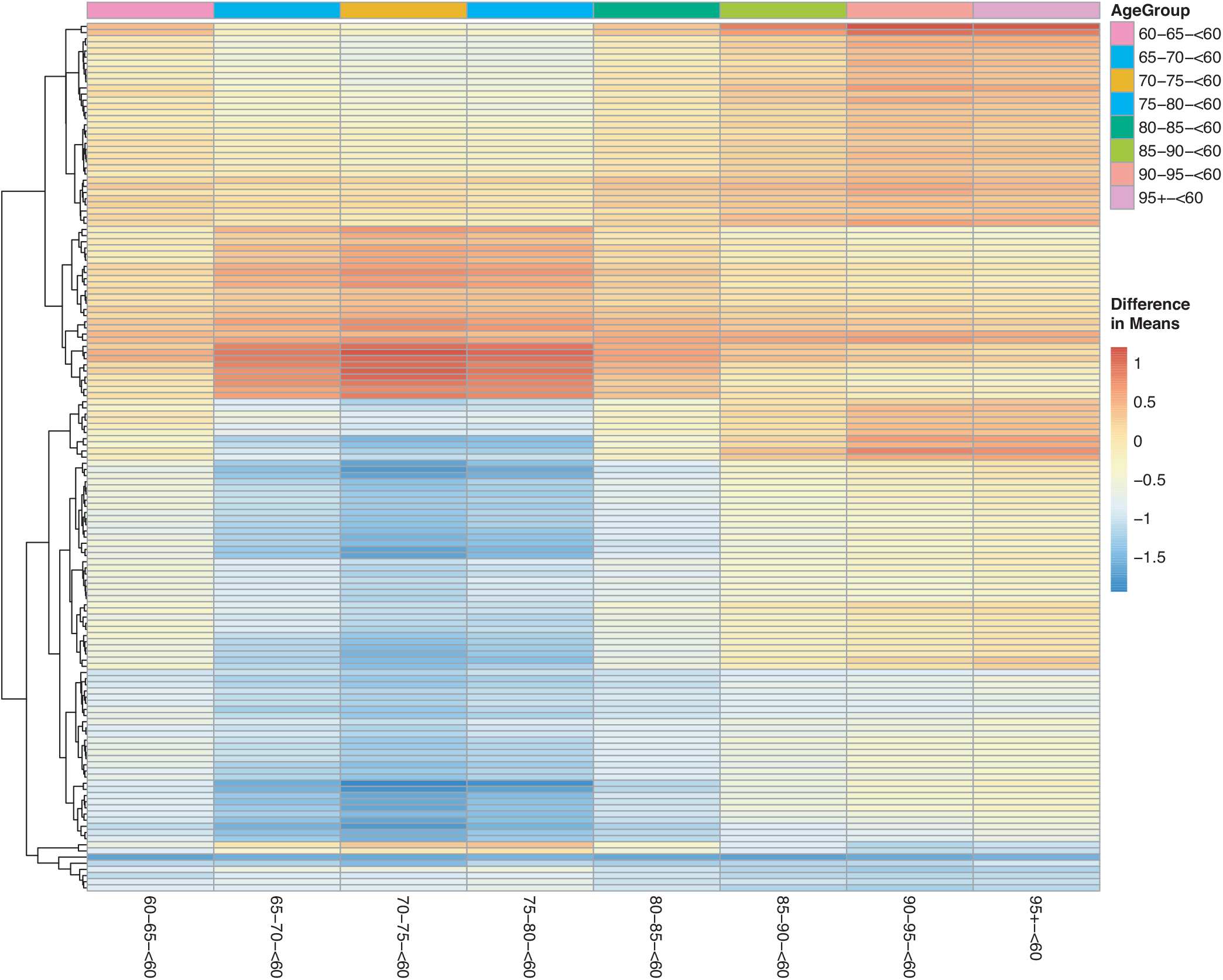
Heatmap with gene clustering to visualize age group effect (difference in means) on the differentially expressed disease (control-AD) gene list.

For tissue effects, we used hippocampus as our baseline due to it being a known target of AD. In addition to filtering for significance, we used again a two-tailed 10% quantile filter ≤−0.6359497 and ≥0.7932871 from the tissue-specific means differences between tissue types (Supplemental Table 1). We found 167 comparisons with tissue differences (see ST11 of online supplemental data) from 125 genes. Our heatmap of these genes show that differences do exist across tissues when compared to hippocampus (Figure 9). For example, nucleus accumbens has higher expression of genes compared to the hippocampus, and putamen has genes that are down-regulated compared to hippocampus (Figure 9). The majority of the expression differences appear to be found in nucleus accumbens and putamen (Figure 9) (see ST11 of online supplemental data). We also assessed how gene expression changes in a given tissue compared to blood (10 %quantile filter: ≤−0.6359497 and ≥0.7932871) (Supplemental Table 1), identifying 152 significant tissue comparisons in 115 genes (see ST12 of online supplemental data). These 115 gene expression profiles across tissues are visualized using the differences of means in Supplemental Figure 9. We again noticed similar trends in the blood comparisons as had in the hippocampus comparisons, with nucleus accumbens showing higher gene expression and putamen lowered expression compared to blood (Supplemental Figure 9).

**Figure 9.**
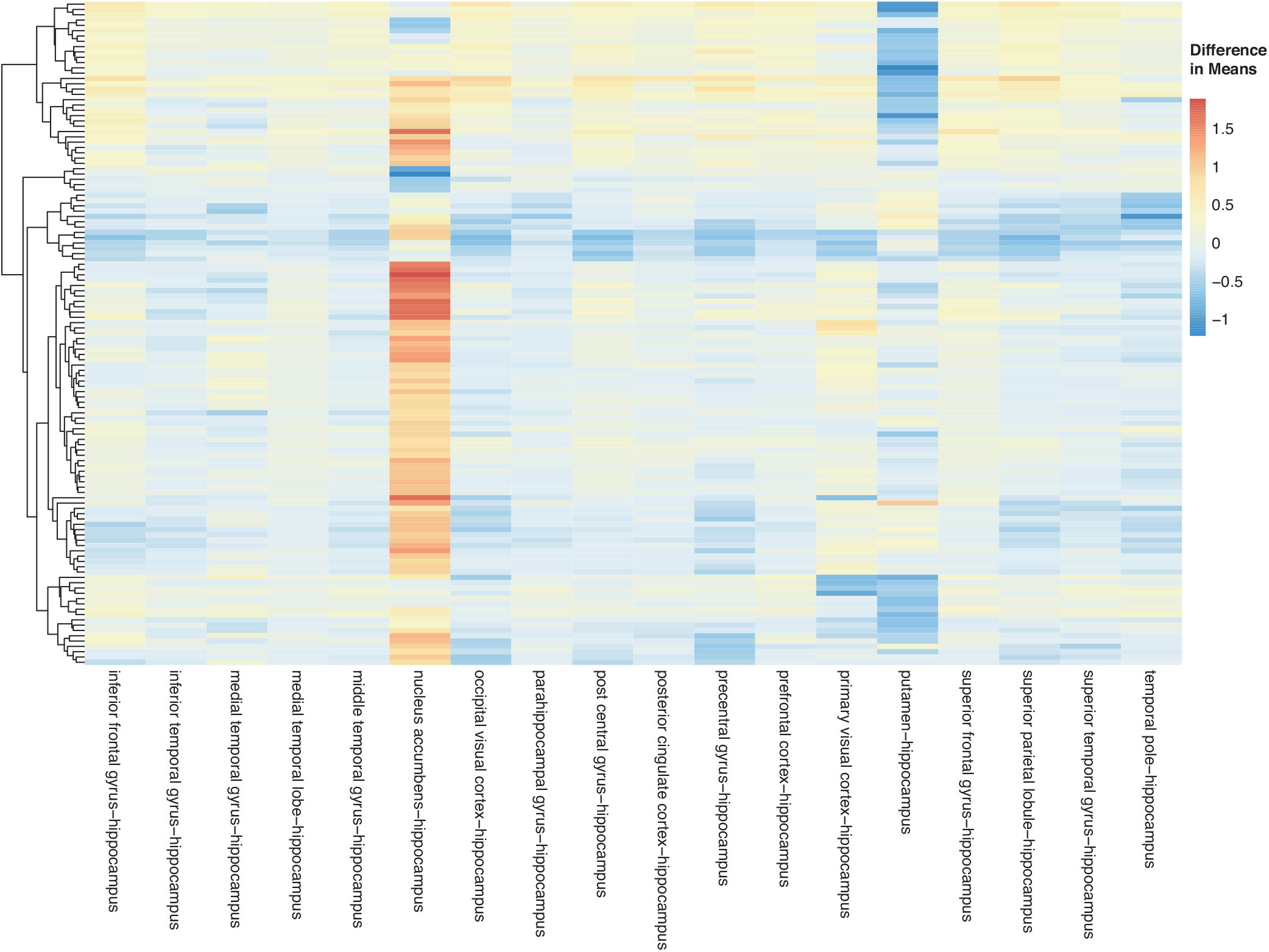
Heatmap with gene clustering to visualize tissue effect (difference in means) on the differentially expressed disease (control-AD) gene list.

## 4 DISCUSSION

As debilitating as Alzheimer’s disease (AD) is, there is still no cure available, and diagnosis is not confidently confirmed until death. There are ongoing research efforts to find biomarkers and gene targets for early detection and intervention in AD. In our study, we investigated changes at the transcript level by conducting a meta-analysis to analyze 8 microarray expression datasets for temporal changes in gene expression due to disease status. In addition to this, we determined if sex, age or tissue type had an effect on gene expression changes in Alzheimer’s associated disease genes. We pre-processed the 8 datasets by background correction, data normalization, and probe annotation. Following this, the datasets were merged into a single dataset (by common gene name) for the meta-analysis. This is the first meta-analysis to explore over 20 different tissues and use a linear model to identify linear and binary effects on gene expression. Our linear model also adjusted batch effects by modeling for the study effect and included age in the model as a linear time series. Modeling with the study factor to account for batch effects was shown to be necessary after exploratory visualization of the expression data before and after combat batch effect correction using principal component analysis to remove variation within the data that was introduced due to different studies(Figures 2,3).

### Significant Gene expression Differences Due to Disease Status and Biological Significance

We first identified significantly different disease genes from ANOVA (see ST4 of online supplemental data), and some of these genes included: APOE,PSEN2,APOD,TREM2,CLU which all have been previously associated with AD. APOE and APOD are members of the apolipoprotein family that transport and metabolize lipids in the central nervous system and play a role in healthy brain function (Elliott et al., 2010). APOE is a strong, well documented, genetic risk factor for AD and polymorphisms in APOE have been shown to affect age of AD onset (Masters et al., 2015). APOD’s mechanism is still not completely understood (Elliott et al., 2010), PSEN2 encodes PS2, an enzyme that cleaves amyloid precursor protein, regulates production of (Aβ), inherited and mutations are associated with early onset (Masters et al., 2015). mutations in CLU lead to lower white matter and increases AD risk ((Braskie et al., 2011; Masters et al., 2015) and TREM2 was identified by GWAS (genome-wide association study) as a disease variant and risk factor for AD (Masters et al., 2015). Our enrichment results of these 3,735 genes were interesting due to them having already been associated with AD in the literature (Table 3, see ST5 and ST6 in online supplemental data). For instance, mitochondrial dysfunction has been previously associated with AD and characterized to cause Aβ deposition, higher production of reactive oxygen species and lowered ATP production (Swerdlow, 2018; Moreira et al., 2010; Onyango et al., 2016). Researchers have also suggested that the immune system plays a role in AD (Heppner et al., 2015; Van Eldik et al., 2016). As for adaptive immune cells, their role in AD is still not clear, however, adaptive immune cells have been shown to reduce AD pathology (Marsh et al., 2016). The loss of B cell production can exacerbate the disease (Marsh et al., 2016). Neurodenegenerative diseases have also been described as having genes that overlap (Moradifard et al., 2018; Wang et al., 2017). Neurodegeneration is closely related to synaptic dysfunction and long term potentiation becomes impaired with age and synaptic dysfunction (Prieto et al., 2017). These results suggest that our meta-analysis is producing disease-related results. Because our KEGG results on the 3,735 genes resulted in identifying the involvement of AD, Parkinson’s and Huntington’s pathways, we investigated if the three neurodegenerative disease signaling pathways had any common genes in our gene list. We determined that AD had 49 genes that overlapped with Huntington’s and 47 with Parkinson’s pathways respectively. We also found that GNAQ, GRIN1 and PLCB1 are in both Huntington’s and AD but not in Parkinson’s pathways, and SNCA is in both Parkinson’s and AD but not Huntington’s pathways. In filtering these genes for biological effect size, PSEN2, APOE, TREM, CLU and other apolipoproteins did not make the cutoff (based on their difference in means between the compared AD/healthy groups). Focusing on the 352 DEG that had a sizable effect size, the down-regulated genes in AD connect with the pathology of the disease (Figure 5. Comparing these 352 DEG to a recently published meta-analysis in which 1400 differentially expressed disease genes were identified, we determined that 136 DEG from our gene list overlapped with theirs, and 216 of our DEG were not in their list (Moradifard et al., 2018). Some stand out unique genes in our list include: GMPR,ABCA1, NOTCH1 and 2, GABRG1,HVCN1,CXCR4, HIP1,MRPS28, FOS. The top up-regulated gene in AD from our meta-analysis, ITPKB, has previously been observed to have over-expression in AD subjects. In a mouse model, the gene was found to be overexpressed and connected to apoptosis, increased (Aβ) production and tau phosphorylation (Stygelbout et al., 2014). CXCR4 (brain development and neuronal cell survival in the hippocampus) (Stelzer et al., 2016), AHNAK (may have a role in development of neuronal cells)(Stelzer et al., 2016), NOTCH1,and NOTCH2 (signaling pathway may be involved in brain development) (Stelzer et al., 2016) were all up-regulated in AD subjects (Table 4. Examples of down-regulared genes: NME1 (neural development) (Stelzer et al., 2016), and mitochondrial proteins MRPL3, MRPS18C (associated with mitchondrial dysfunction observed in AD) were down-regulated in AD samples 4.

### Sex, Age and Tissue Effect on Disease Status Biologically Significant Genes

For the sex factor, we determined that 46 of our DEG (23 up- and down- regulated in males compared to females) had a sex effect. Figure 7 highlights the enriched pathways from up-regulated genes in males. This indicates that pathways important for neuronal system and chemical synapses were down-regulated in females (Figure 7). This is also supported by the current literature, which indicates that women are at higher risk for AD (Vina and Lloret, 2010; Podcasy and Epperson, 2016; Seshadri et al., 1997). This increased risk by sex is due to the loss of estrogen protection (due to menopause) against (Aβ)’s toxicity on the mitochondria (Vina and Lloret, 2010; Podcasy and Epperson, 2016). Older women produce more reactive oxygen species with the decline in estrogen levels (Vina and Lloret, 2010; Podcasy and Epperson, 2016). Estrogen replacement therapy is a treatment for AD, and it is being determined that estrogen works by increasing the expression of antioxidant genes (Vina and Lloret, 2010; Podcasy and Epperson, 2016). A recently published meta-analysis also explored sex effects on AD gene expression (Moradifard et al., 2018). Moradifard et al., found male and female specific AD associated genes and genes that overlapped in both sexes (Moradifard et al., 2018). Of the 46 disease associated genes we found to be affected by sex, 22 were found in both males and females, 9 only in males, and 5 only in females in Moradifard et al gene list. 10 of our sex impacted disease genes (CYBRD1,DIRAS2,FAM107B,FOS,GMPR,HVCN1,ITIH5,MAPK,RNF135,SLC40A1) did not overlap with their findings, and these genes have been previously associated with oxidative stress, cell signaling and transport, apoptosis and AD. For instance, GMPR was found to gradually increase as AD progressed (Liu et al., 2018). It produces GMPR1 which is associated with the phosphorylation of tau (Liu et al., 2018).

Aging trends on the diseased genes were visualized in Figure 8. Subjects <60 were used as a baseline because on average, AD symptoms start at ages 65 and older. We observed clear age-related patterns when looking at the difference of means between age cohorts for the disease gene list, Figure 8 (see ST10 of online supplemental data). Highlighting a few of the changes: SNAP9 which is involved in synaptic transmission and associated with late onset (Zhang et al., 2013), STMN2 which is necessary for microtubule dynamics and neuronal growth (Antonsson et al., 1998; Chiellini et al., 2008), and SST, a neuropeptide that interacts with (Aβ) and can influence how it aggregates (Solarski et al., 2018; Hama and Saido, 2005) were all up-regulated in <60 age group(see ST8 of online supplemental data). Also, STMN2 and SST have both previously been associated with expression reduction due to age(Solarski et al., 2018; Stelzer et al., 2016).

ABCA1,GMPR, HVCN1, ITPKB, NOTCH1 all had higher expression in older age groups compared to the baseline. Our findings highlight genes previously associated with AD and their temporal trends and also some additional genes that experience age-effects (see Figure 8, and ST10 of online supplemental data)

For investigating tissue effects, we used hippocampus (232 samples) as a baseline due to its having being identified as one of the first regions to be affected by AD (Masters et al., 2015). We also used blood (519 samples) as a baseline to explore an underdeveloped non-invasive approach to monitoring AD. In both analyses, we see similar trends with nucleus accumbens (51 samples) and putamen (52 samples) showing greater differences in expression (Figure 9 and Supplemental Figure 9). The distribution of samples per tissue type was inconsistent with hippocampus and blood having larger number of samples compared to an average of around 55 samples per tissue in other categories. These results show the potential of blood and other tissues for monitoring gene expression changes in AD, but also the need for further focused mechanistic studies in different tissues.

### Limitations of the Study

Using publicly available data introduced limitations to our research design. Lack of uniform annotation and missing information across datasets can make conducting a meta-analysis on multiple datasets challenging. The number of datasets used in our meta-analysis was limited by poor annotations that could not meet our selection criteria, and this in turn placed bounds to our sample size and power of the study. Our analysis was also unbalanced: 2,088 samples made up of 771 healthy controls, 868 AD subjects, 449 subjects reported as possibly having AD, 1308 females and 780 males, and the breakdown of age groups is also somewhat uneven. The available public data used for our meta-analysis also lacked diversity in samples, because in most datasets race and ethnicity are not reported. This information would be helpful particularly since AD has been reported by the CDC to be more prevalent in African Americans (Centers for Disease Control and Prevention, 2018; Steenland et al., 2016). In addition, the use of micro-array expression data for meta-analysis is a limitation in terms of not being able to query the entire transcriptome or query novel genes. Also, in our merged dataset, large variability was introduced in data due to the large number of tissues (26) and methods used for extractions (study effect), which we attempted to correct for by utilizing both as factors in our model, and including binary interaction terms as well.

### Future Directions and Recommendations

Our study provides gene lists by factor (disease status, sex, age and tissue) of differentially expressed genes. To expand on this research, the use of RNA-sequencing data can reveal novel differentially expressed genes, biomarkers and gene targets for AD. In addition to RNA-sequencing, implementing other omics technologies such as proteomics and metabolomics can help to fully describe the pathology of AD, and identify additional biomarkers for early detection. Including data with racial diversity is also necessary. AD has higher prevalence in African Americans (Steenland et al., 2016). Due to reports of racial differences in AD, with an AD prevalence breakdown of: 14% of African American population compared to 12% in Hispanics and 10% in whites (Centers for Disease Control and Prevention, 2018), including racial diversity in future studies would help identify this potential variability in susceptibility and identify if certain treatments might be better suited in some races than others. Improving the representation of races in clinical trials and molecular reports of AD can help with health disparities within the field. Exploring the use of easily accessible tissues, such as blood, to monitor changes in target genes/biomarkers might also prove helpful for early detection and provide a more systems-level understanding of AD. Determining the best or novel biomarkers to track for AD requires exploring also mechanistic aspects of the disease. For example, monitoring exosomes and autoantibodies which can be connected to the dysfunction of the immune system is one mode of action that is being associated with AD (O’Bryant, 2016). Lastly, as omics technologies advance, implementing personalized omics for early detection and treatment may prove useful in improving individual AD outcomes with the increase in the aging population.

## Supporting information

## AUTHOR CONTRIBUTIONS

LB and GM wrote the manuscript.

## ACKNOWLEDGMENTS

An early version of this work was presented at the 2018 American Society of Human Genetics Annual Conference in San Diego abstract # 1369T.

## SUPPLEMENTAL DATA

Our supplemental figures and tables are provided in the accompanying supplementary data pdf file. All of our datasets/results from our meta-analysis pipeline have been uploaded to FigShare as online supplemental data, as described below (the corresponding online file names begin with a prefix ‘ST’ and are enumerated as also referred to in the manuscript).

## DATA AVAILABILITY STATEMENT

The datasets generated and analyzed for this study can be found in Figshare’s online digital repository at https://doi.org/10.6084/m9.figshare.7435469.

## REFERENCES

Antonsson, B., Kassel, D. B., Di Paolo, G., Lutjens, R., Riederer, B. M., and Grenningloh, G. (1998). Identification of in vitro phosphorylation sites in the growth cone protein scg10. effect of phosphorylation site mutants on microtubule-destabilizing activity. J Biol Chem 273, 8439–46

Berchtold, N. C., Cribbs, D. H., Coleman, P. D., Rogers, J., Head, E., Kim, R., et al. (2008). Gene expression changes in the course of normal brain aging are sexually dimorphic. Proceedings of the National Academy of Sciences

Black, S., De Gregorio, E., and Rappuoli, R. (2015). Developing vaccines for an aging population. Science translational medicine 7, 281ps8–281ps8. doi:10.1126/scitranslmed.aaa0722

Blalock, E. M., Buechel, H. M., Popovic, J., Geddes, J. W., and Landfield, P. W. (2011). Microarray analyses of laser-captured hippocampus reveal distinct gray and white matter signatures associated with incipient alzheimer’s disease. Journal of chemical neuroanatomy 42, 118–126

Bland, J. M. and Altman, D. G. (1995). Multiple significance tests: the bonferroni method. BMJ 310, 170. doi:10.1136/bmj.310.6973.170

Braskie, M. N., Jahanshad, N., Stein, J. L., Barysheva, M., McMahon, K. L., de Zubicaray, G. I., et al. (2011). Common alzheimer’s disease risk variant within the clu gene affects white matter microstructure in young adults. The Journal of neuroscience : the official journal of the Society for Neuroscience 31, 6764–6770. doi:10.1523/JNEUROSCI.5794-10.2011

Brazma, A., Parkinson, H., Sarkans, U., Shojatalab, M., Vilo, J., Abeygunawardena, N., et al. (2003). Arrayexpress—a public repository for microarray gene expression data at the ebi. Nucleic acids research 31, 68–71. doi:https://doi.org/10.1093/nar/gkg091

Bronner, I. F., Bochdanovits, Z., Rizzu, P., Kamphorst, W., Ravid, R., van Swieten, J. C., et al. (2009). Comprehensive mrna expression profiling distinguishes tauopathies and identifies shared molecular pathways. PLoS One 4, e6826

Brookmeyer, R., Abdalla, N., Kawas, C. H., and Corrada, M. M. (2018). Forecasting the prevalence of preclinical and clinical alzheimer’s disease in the united states. Alzheimers Dement 14, 121–129. doi:10.1016/j.jalz.2017.10.009

[Dataset] Centers for Disease Control and Prevention (2018). U.s. burden of alzheimer’s disease, related dementias to double by 2060. Accessed: 2018-12-01

Chiellini, C., Grenningloh, G., Cochet, O., Scheideler, M., Trajanoski, Z., Ailhaud, G., et al. (2008). Stathmin-like 2, a developmentally-associated neuronal marker, is expressed and modulated during osteogenesis of human mesenchymal stem cells. Biochem Biophys Res Commun 374, 64–8. doi:10.1016/j.bbrc.2008.06.121

Childs, B. G., Durik, M., Baker, D. J., and Van Deursen, J. M. (2015). Cellular senescence in aging and age-related disease: from mechanisms to therapy. Nature medicine 21, 1424

Dallmeyer, S., Wicker, P., and Breuer, C. (2017). How an aging society affects the economic costs of inactivity in germany: empirical evidence and projections. European Review of Aging and Physical Activity 14, 18

De Jager, P., Ma, Y., McCabe, C., Xu, J., Vardarajan, B. N., Felsky, D., et al. (2018). A multi-omic atlas of the human frontal cortex for aging and alzheimer’s disease research. bioRxiv, 251967

Drayer, B. P. (1988). Imaging of the aging brain. Radiology 166, 785–796

Edgar, R., Domrachev, M., and Lash, A. E. (2002). Gene expression omnibus: Ncbi gene expression and hybridization array data repository. Nucleic Acids Res 30, 207–10. doi:https://doi.org/10.1093/nar/30.1.207

Elliott, D. A., Weickert, C. S., and Garner, B. (2010). Apolipoproteins in the brain: implications for neurological and psychiatric disorders. Clinical lipidology 51, 555–573. doi:10.2217/CLP.10.37

Hama, E. and Saido, T. C. (2005). Etiology of sporadic alzheimer’s disease: somatostatin, neprilysin, and amyloid beta peptide. Med Hypotheses 65, 498–500. doi:10.1016/j.mehy.2005.02.045

Hebert, L. E., Weuve, J., Scherr, P. A., and Evans, D. A. (2013). Alzheimer disease in the united states (2010-2050) estimated using the 2010 census. Neurology 80, 1778–83. doi:10.1212/WNL.0b013e31828726f5

Heppner, F. L., Ransohoff, R. M., and Becher, B. (2015). Immune attack: the role of inflammation in alzheimer disease. Nat Rev Neurosci 16, 358–72. doi:10.1038/nrn3880

[Dataset] Irizarry, R. and Love, M. (2015). Ph525x series - biomedical data science. Accessed: 2018-01-18

Jaul, E. and Barron, J. (2017). Age-related diseases and clinical and public health implications for the 85 years old and over population. Front Public Health 5, 335. doi:10.3389/fpubh.2017.00335

Jevtic, S., Sengar, A. S., Salter, M. W., and McLaurin, J. (2017). The role of the immune system in alzheimer disease: Etiology and treatment. Ageing Res Rev 40, 84–94. doi:10.1016/j.arr.2017.08.005

Johnson, W. E., Li, C., and Rabinovic, A. (2007). Adjusting batch effects in microarray expression data using empirical bayes methods. Biostatistics 8, 118–127

Kanehisa, M., Furumichi, M., Tanabe, M., Sato, Y., and Morishima, K. (2017). Kegg: new perspectives on genomes, pathways, diseases and drugs. Nucleic Acids Res 45, D353–D361. doi:10.1093/nar/gkw1092

Kanehisa, M. and Goto, S. (2000). Kegg: kyoto encyclopedia of genes and genomes. Nucleic Acids Res 28, 27–30. doi:https://doi.org/10.1093/nar/28.1.27

Kanehisa, M., Sato, Y., Kawashima, M., Furumichi, M., and Tanabe, M. (2016). Kegg as a reference resource for gene and protein annotation. Nucleic Acids Res 44, D457–62. doi:10.1093/nar/gkv1070

Liang, W. S., Dunckley, T., Beach, T. G., Grover, A., Mastroeni, D., Walker, D. G., et al. (2007). Gene expression profiles in anatomically and functionally distinct regions of the normal aged human brain. Physiological genomics 28, 311–322

Liu, H., Luo, K., and Luo, D. (2018). Guanosine monophosphate reductase 1 is a potential therapeutic target for alzheimer’s disease. Scientific Reports 8, 2759. doi:10.1038/s41598-018-21256-6

Lopez-Otin, C., Blasco, M. A., Partridge, L., Serrano, M., and Kroemer, G. (2013). The hallmarks of aging. Cell 153, 1194–217. doi:10.1016/j.cell.2013.05.039

Maere, S., Heymans, K., and Kuiper, M. (2005). Bingo: a cytoscape plugin to assess overrepresentation of gene ontology categories in biological networks. Bioinformatics 21, 3448–9. doi:10.1093/bioinformatics/bti551

Marsh, S. E., Abud, E. M., Lakatos, A., Karimzadeh, A., Yeung, S. T., Davtyan, H., et al. (2016). The adaptive immune system restrains alzheimer’s disease pathogenesis by modulating microglial function. Proc Natl Acad Sci U S A 113, E1316–25. doi:10.1073/pnas.1525466113

Masters, C. L., Bateman, R., Blennow, K., Rowe, C. C., Sperling, R. A., and Cummings, J. L. (2015). Alzheimer’s disease. Nature Reviews Disease Primers 1, 15056. doi:10.1038/nrdp.2015.56

Matthews, K. A., Xu, W., Gaglioti, A. H., Holt, J. B., Croft, J. B., Mack, D., et al. (2018). Racial and ethnic estimates of alzheimer’s disease and related dementias in the united states (2015-2060) in adults aged > 65 years. Alzheimer’s & Dementia doi:https://doi.org/10.1016/j.jalz.2018.06.3063

Mattson, M. P. and Arumugam, T. V. (2018). Hallmarks of brain aging: Adaptive and pathological modification by metabolic states. Cell Metab 27, 1176–1199. doi:10.1016/j.cmet.2018.05.011

Mias, G. (2018a). Analysis of Variance for Multiple Tests (Cham: Springer International Publishing), chap. 6.3. 133–170. doi:10.1007/978-3-319-72377-8_4

Mias, G. (2018b). Databases: E-Utilities and UCSC Genome Browser (Cham: Springer International Publishing), chap. 4. 133–170. doi:10.1007/978-3-319-72377-8_4

Mias, G. I., Yusufaly, T., Roushangar, R., Brooks, L. R., Singh, V. V., and Christou, C. (2016). Mathiomica: An integrative platform for dynamic omics. Sci Rep 6, 37237. doi:10.1038/srep37237

Miller, J. A., Woltjer, R. L., Goodenbour, J. M., Horvath, S., and Geschwind, D. H. (2013). Genes and pathways underlying regional and cell type changes in alzheimer’s disease. Genome medicine 5, 48

Moradifard, S., Hoseinbeyki, M., Ganji, S. M., and Minuchehr, Z. (2018). Analysis of microrna and gene expression profiles in alzheimer’s disease: A meta-analysis approach. Scientific reports 8, 4767

Moreira, P. I., Carvalho, C., Zhu, X., Smith, M. A., and Perry, G. (2010). Mitochondrial dysfunction is a trigger of alzheimer’s disease pathophysiology. Biochimica et Biophysica Acta (BBA)-Molecular Basis of Disease 1802, 2–10. doi:https://doi.org/10.1016/j.bbadis.2009.10.006

Nygaard, V., Rødland, E. A., and Hovig, E. (2016). Methods that remove batch effects while retaining group differences may lead to exaggerated confidence in downstream analyses. Biostatistics 17, 29–39

O’Bryant, S. E. (2016). Introduction to special issue on advances in blood-based biomarkers of alzheimer’s disease. Alzheimer’s & dementia (Amsterdam, Netherlands) 3, 110–112. doi:10.1016/j.dadm.2016.06.003

Onyango, I. G., Dennis, J., and Khan, S. M. (2016). Mitochondrial dysfunction in alzheimer’s disease and the rationale for bioenergetics based therapies. Aging Dis 7, 201–14. doi:10.14336/AD.2015.1007

Pavlidis, P. (2003). Using anova for gene selection from microarray studies of the nervous system. Methods 31, 282–289

Podcasy, J. L. and Epperson, C. N. (2016). Considering sex and gender in alzheimer disease and other dementias. Dialogues Clin Neurosci 18, 437–446

Prieto, G. A., Trieu, B. H., Dang, C. T., Bilousova, T., Gylys, K. H., Berchtold, N. C., et al. (2017). Pharmacological rescue of long-term potentiation in alzheimer diseased synapses. J Neurosci 37, 1197–1212. doi:10.1523/JNEURoSCI.2774-16.2016

R Core Team (2018). R: A Language and Environment for Statistical Computing. R Foundation for Statistical Computing, Vienna, Austria

Rowe, J. W., Fulmer, T., and Fried, L. (2016). Preparing for better health and health care for an aging population. JAMA 316, 1643–1644. doi:10.1001/jama.2016.12335

Sakia, R. (1992). The box-cox transformation technique: a review. The statistician, 169–178

Seshadri, S., Wolf, P. A., Beiser, A., Au, R., McNulty, K., White, R., et al. (1997). Lifetime risk of dementia and alzheimer’s disease. the impact of mortality on risk estimates in the framingham study. Neurology 49, 1498–504

Shannon, P., Markiel, A., Ozier, O., Baliga, N. S., Wang, J. T., Ramage, D., et al. (2003). Cytoscape: a software environment for integrated models of biomolecular interaction networks. Genome Res 13, 2498–504. doi:10.1101/gr.1239303

Solarski, M., Wang, H., Wille, H., and Schmitt-Ulms, G. (2018). Somatostatin in alzheimer’s disease: A new role for an old player. Prion 12, 1–8. doi:10.1080/19336896.2017.1405207

Sood, S., Gallagher, I. J., Lunnon, K., Rullman, E., Keohane, A., Crossland, H., et al. (2015). A novel multi-tissue rna diagnostic of healthy ageing relates to cognitive health status. Genome biology 16, 185

Steenland, K., Goldstein, F. C., Levey, A., and Wharton, W. (2016). A meta-analysis of alzheimer’s disease incidence and prevalence comparing african-americans and caucasians. Journal of Alzheimer’s disease : JAD 50, 71–76. doi:10.3233/JAD-150778

Stelzer, G., Rosen, N., Plaschkes, I., Zimmerman, S., Twik, M., Fishilevich, S., et al. (2016). The genecards suite: From gene data mining to disease genome sequence analyses. Curr Protoc Bioinformatics 54, 1 30 1–1 30 33. doi:10.1002/cpbi.5

Stygelbout, V., Leroy, K., Pouillon, V., Ando, K., D’amico, E., Jia, Y., et al. (2014). Inositol trisphosphate 3-kinase b is increased in human alzheimer brain and exacerbates mouse alzheimer pathology. Brain 137, 537–552. doi:10.1093/brain/awt344

Swerdlow, R. H. (2018). Mitochondria and mitochondrial cascades in alzheimer’s disease. J Alzheimers Dis 62, 1403–1416. doi:10.3233/JAD-170585

Taylor, C. A., Greenlund, S. F., McGuire, L. C., Lu, H., and Croft, J. B. (2017). Deaths from alzheimer’s disease-united states, 1999-2014. MMWR. Morbidity and mortality weekly report 66, 521

Toepper, M. (2017). Dissociating normal aging from alzheimer’s disease: A view from cognitive neuroscience. J Alzheimers Dis 57, 331–352. doi:10.3233/JAD-161099

Tukey, J. W. (1949). Comparing individual means in the analysis of variance. Biometrics 5, 99–114. doi:10.2307/3001913

[Dataset] United Nations Department of Economic and Social Affairs (2015). World population ageing 2015. Accessed: 2018-11-01

Van Deursen, J. M. (2014). The role of senescent cells in ageing. Nature 509, 439

Van Eldik, L. J., Carrillo, M. C., Cole, P. E., Feuerbach, D., Greenberg, B. D., Hendrix, J. A., et al. (2016). The roles of inflammation and immune mechanisms in alzheimer’s disease. Alzheimer’s & Dementia: Translational Research & Clinical Interventions 2, 99–109. doi:10.1016/j.trci.2016.05.001

Vina, J. and Lloret, A. (2010). Why women have more alzheimer’s disease than men: gender and mitochondrial toxicity of amyloid-β peptide. Journal of Alzheimer’s disease 20, S527–S533. doi: 10.3233/JAD-2010-100501

Wang, M., Roussos, P., McKenzie, A., Zhou, X., Kajiwara, Y., Brennand, K. J., et al. (2016). Integrative network analysis of nineteen brain regions identifies molecular signatures and networks underlying selective regional vulnerability to alzheimer’s disease. Genome Med 8, 104. doi:10.1186/s13073-016-0355-3

Wang, Q., Li, W. X., Dai, S. X., Guo, Y. C., Han, F. F., Zheng, J. J., et al. (2017). Meta-analysis of parkinson’s disease and alzheimer’s disease revealed commonly impaired pathways and dysregulation of nrf2-dependent genes. J Alzheimers Dis 56, 1525–1539. doi:10.3233/JAD-161032

Winkler, J. M. and Fox, H. S. (2013). Transcriptome meta-analysis reveals a central role for sex steroids in the degeneration of hippocampal neurons in alzheimer’s disease. BMC Syst Biol 7, 51. doi:10.1186/1752-0509-7-51

Wolfram Research, Inc. (2017). Mathematica, version 11.2 edn.

Yu, G. and He, Q. Y. (2016). Reactomepa: an r/bioconductor package for reactome pathway analysis and visualization. Mol Biosyst 12, 477–9. doi:10.1039/c5mb00663e

Yu, G., Wang, L. G., Han, Y., and He, Q. Y. (2012). clusterprofiler: an r package for comparing biological themes among gene clusters. OMICS 16, 284–7. doi:10.1089/omi.2011.0118

Zhang, B., Gaiteri, C., Bodea, L. G., Wang, Z., McElwee, J., Podtelezhnikov, A. A., et al. (2013). Integrated systems approach identifies genetic nodes and networks in late-onset alzheimer’s disease. Cell 153, 707–20. doi:10.1016/j.cell.2013.03.030

