## Supplementary material for "Data-Driven Analysis of Age, Sex, and Tissue Effects on Gene Expression Variability in Alzheimer’s Disease"

### 1 SUPPLEMENTARY TABLES AND FIGURES

#### 1.1 Tables

| Quantile | Disease (control-AD) | Sex (male-female) | AgeGroup (i-<60) | Tissue (i-blood) | Tissue (i-hippocamppus) |
|---|---|---|---|---|---|
| 0.1% | -0.1980181 | -0.1850368 | -1.8342734 | -1.4193659 | -1.2176260 |
| 1% | -0.1409218 | -0.1662240 | -1.5978577 | -1.1791001 | -0.8806144 |
| 2.5% | -0.1242022 | -0.1531487 | -1.4093136 | -0.9535290 | -0.6994610 |
| 5% | -0.1109185 | -0.1286464 | -1.2380410 | -0.7905619 | -0.5988491 |
| **10%** | **-0.0944796** | **-0.0863796** | **-1.0477827** | **-0.6359497** | **-0.5187091** |
| **90%** | **0.1195751** | **0.2502144** | **0.3308682** | **0.7932871** | **0.8181017** |
| 95% | 0.1398357 | 0.2678312 | 0.4815650 | 1.0342074 | 1.0113406 |
| 97.5% | 0.1597702 | 0.2788782 | 0.6502154 | 1.2459840 | 1.2049823 |
| 99% | 0.1851621 | 0.3036726 | 0.8852537 | 1.5229805 | 1.6578388 |
| 99.9% | 0.2625072 | 0.3698125 | 1.1441852 | 1.7230551 | 1.7531744 |

**Table S1.** Quantiles on differences of means between group comparisons from TukeyHSD analysis for each factor with the 10% and 90% highlighted.

| Gene | diff | lwr | upr | tukey.p.adj |
|---|---|---|---|---|
| SNAP91 | 0.38789336 | 0.3162492 | 0.45953751 | <5.91E-12 |
| AMPH | 0.261109 | 0.227946 | 0.29427199 | 5.91E-12 |
| CABP1 | 0.25221566 | 0.2197738 | 0.28465749 | 5.91E-12 |
| CCK | 0.2736884 | 0.2290262 | 0.31835057 | 5.91E-12 |
| CHGB | 0.27223361 | 0.2330743 | 0.31139295 | 5.91E-12 |
| CPQ | -0.15134075 | -0.1739617 | -0.12871979 | 5.91E-12 |
| CXCR4 | -0.18569224 | -0.2153576 | -0.15602687 | 5.91E-12 |
| DIRAS2 | 0.26469562 | 0.2210822 | 0.30830901 | 5.91E-12 |
| EEF1A2 | 0.29830496 | 0.2546981 | 0.34191185 | 5.91E-12 |
| GABRG2 | 0.28727303 | 0.2421887 | 0.33235734 | 5.91E-12 |
| GFAP | 0.26148848 | 0.2372962 | 0.28568071 | 5.91E-12 |
| GJA1 | 0.30536761 | 0.279178 | 0.33155722 | 5.91E-12 |
| KLF2 | -0.16010858 | -0.1822269 | -0.13799028 | 5.91E-12 |
| MYT1L | 0.25975404 | 0.2232677 | 0.29624039 | 5.91E-12 |
| NEFL | 0.27515335 | 0.2353672 | 0.31493946 | 5.91E-12 |
| NRN1 | 0.26422817 | 0.2300824 | 0.29837391 | 5.91E-12 |
| RGS4 | 0.27860758 | 0.2432385 | 0.31397667 | 5.91E-12 |
| SERPINI1 | 0.26217204 | 0.2301731 | 0.29417102 | 5.91E-12 |
| SH3GL2 | 0.30717515 | 0.2681965 | 0.34615382 | 5.91E-12 |
| IL13RA1 | -0.15586061 | -0.1816785 | -0.13004272 | 5.91E-12 |
| ERC2 | 0.26822985 | 0.2230493 | 0.31341042 | 5.91E-12 |
| GAD1 | 0.26177649 | 0.2178336 | 0.30571939 | 5.91E-12 |
| SLC40A1 | -0.18276614 | -0.2136628 | -0.15186946 | 5.91E-12 |
| ITIH5 | -0.16639611 | -0.1947932 | -0.137999 | 5.91E-12 |
| FAM19A1 | 0.269102 | 0.2230286 | 0.31517537 | 5.91E-12 |
| FGF13 | 0.25310759 | 0.2088214 | 0.29739382 | 5.92E-12 |
| AHNAK | -0.10311728 | -0.1242383 | -0.08199628 | 5.93E-12 |
| RPA3 | -0.13337106 | -0.1607507 | -0.10599143 | 5.93E-12 |
| EZR | -0.1182311 | -0.141882 | -0.09458025 | 5.93E-12 |
| ITPKB | -0.11649742 | -0.1417882 | -0.09120668 | 5.93E-12 |
| GABRA1 | 0.27928408 | 0.2277225 | 0.33084567 | 5.93E-12 |
| MAP3K1 | -0.16567888 | -0.1964814 | -0.13487641 | 5.93E-12 |
| NOTCH1 | -0.10639043 | -0.1270303 | -0.0857506 | 5.93E-12 |
| HVCN1 | -0.10966873 | -0.1333269 | -0.0860106 | 5.93E-12 |
| PCDH8 | 0.26623656 | 0.2038808 | 0.32859232 | 5.93E-12 |
| LDLRAP1 | -0.13026125 | -0.1611422 | -0.09938027 | 5.93E-12 |
| GMPR | -0.14621751 | -0.1812927 | -0.11114232 | 5.94E-12 |
| CYBRD1 | -0.1288122 | -0.1605469 | -0.09707747 | 5.94E-12 |
| PRKD2 | -0.09889483 | -0.1237886 | -0.07400105 | 5.94E-12 |
| PRKX | -0.12798343 | -0.1609683 | -0.09499854 | 5.97E-12 |
| STMN2 | 0.25735011 | 0.1839117 | 0.33078852 | 8.42E-12 |
| HIP1 | -0.11198115 | -0.1436469 | -0.08031542 | 1.16E-11 |
| FOS | -0.15132275 | -0.1952695 | -0.10737604 | 2.55E-11 |
| FAM107B | -0.10351231 | -0.1344653 | -0.07255936 | 7.72E-11 |
| RNF135 | -0.0875083 | -0.1183122 | -0.05670445 | 2.92E-08 |
| ID3 | -0.10925012 | -0.1502501 | -0.06825012 | 1.94E-07 |

**Table S2.** TukeyHSD results table of statistically significant differentially expressed disease genes with sex effect.

### 1.2 Figures

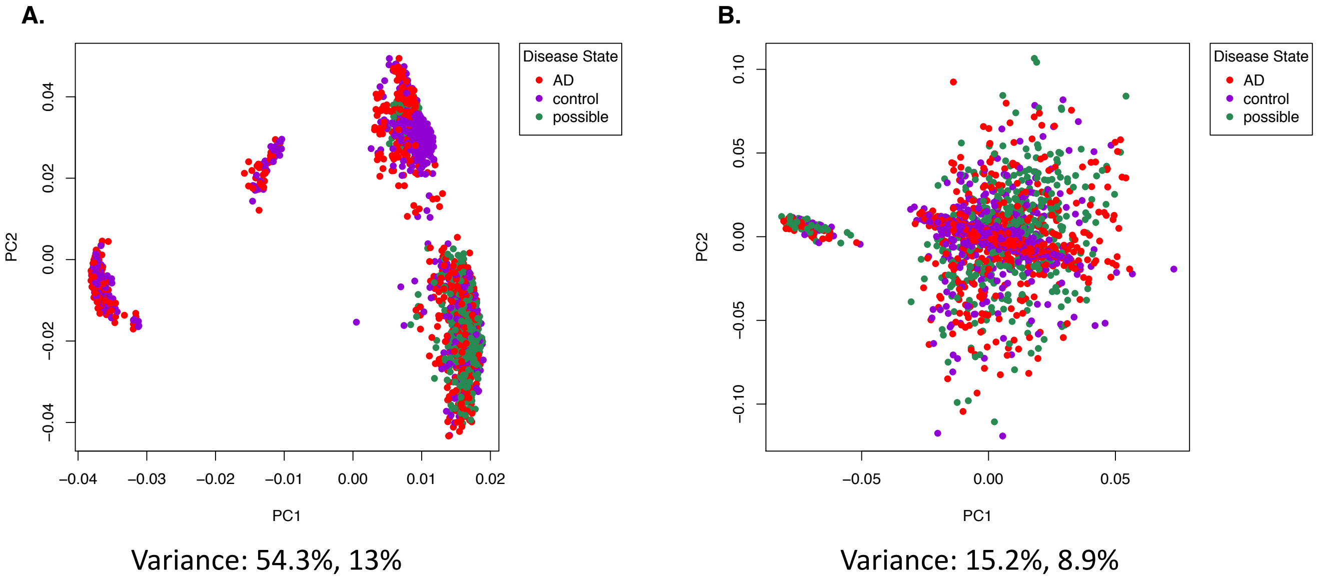


**Figure S1.** Principal component analysis of the disease factor before and after batch correction with ComBat

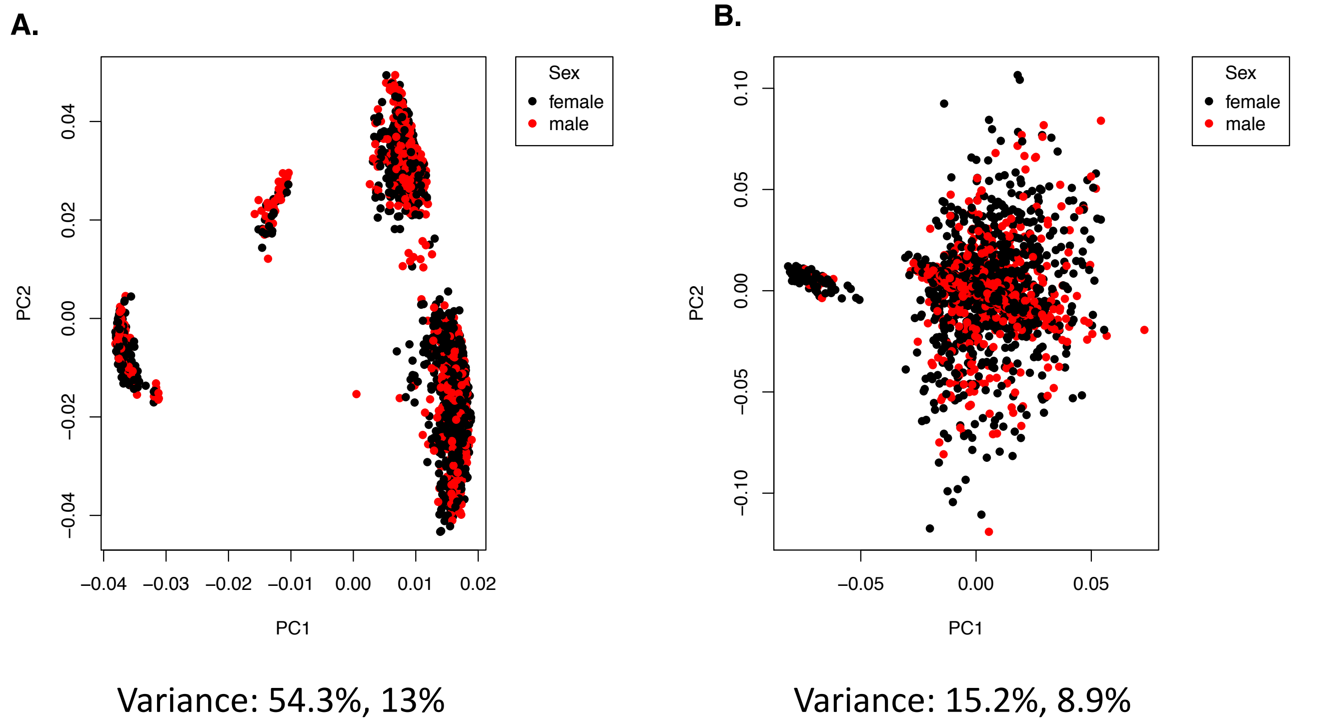


**Figure S2.** Principal component analysis of the sex factor before and after batch effect correction with ComBat.

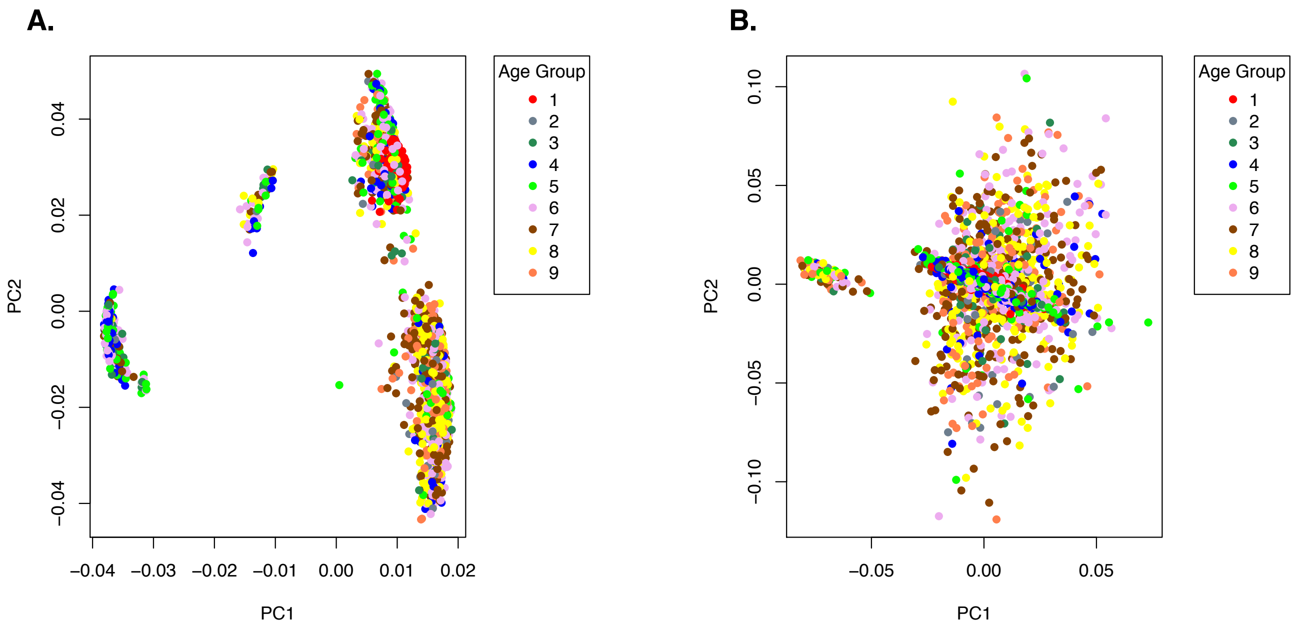


Variance: 54.3%, 13%

Variance: 15.2%, 8.9%

**Figure S3.** Principal component analysis of the age group factor before and after batch effect correction with ComBat.

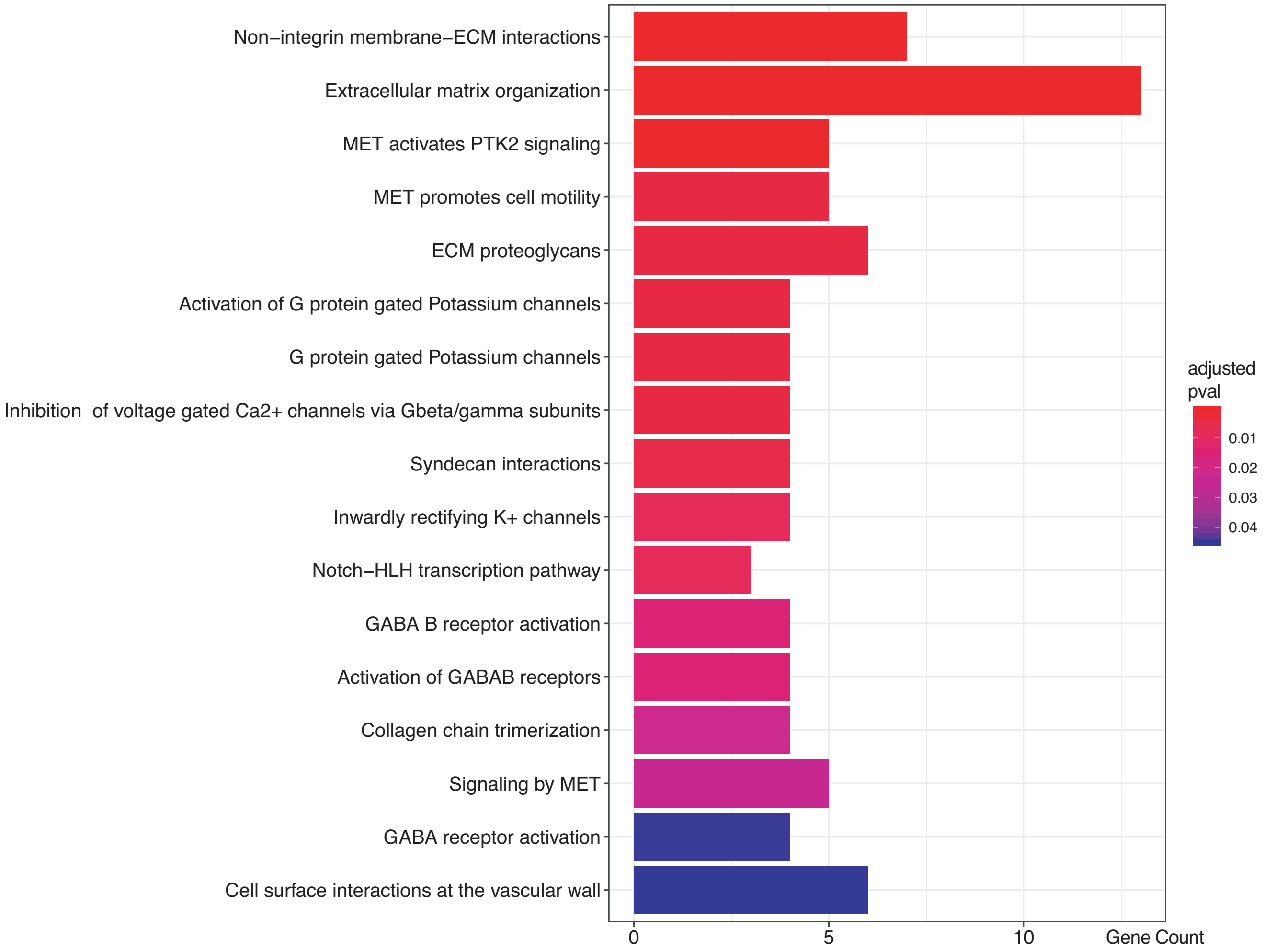


**Figure S4.** Reactome pathway analysis bar plot of enriched pathways and number of gene hits. Gene list: Genes that are up-regulated in Alzheimer's disease but down-regulated in healthy controls.

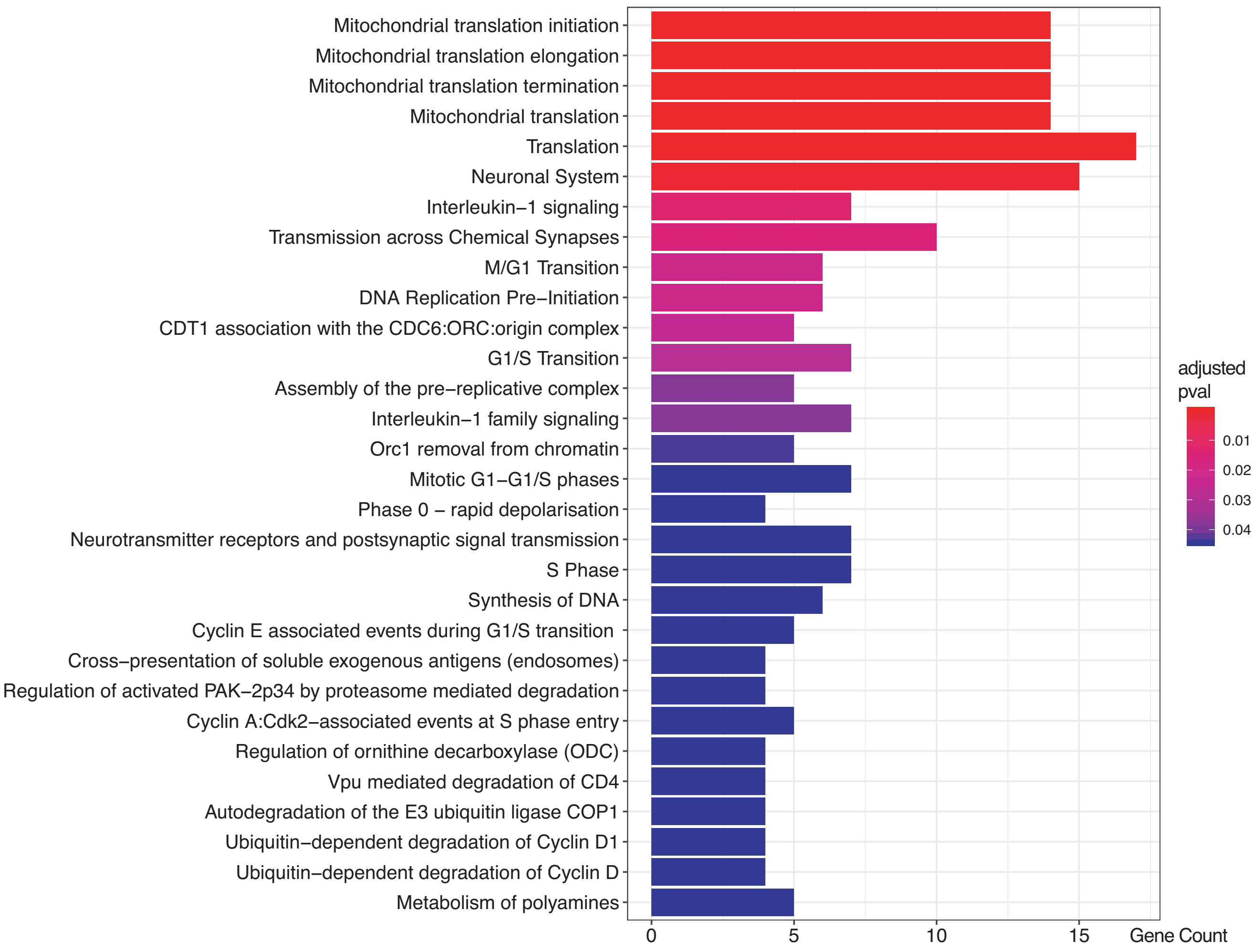


**Figure S5.** Reactome pathway analysis bar plot of enriched pathways and number of gene hits. Gene list: Genes that are down-regulated in Alzheimer's disease but up-regulated in healthy controls.

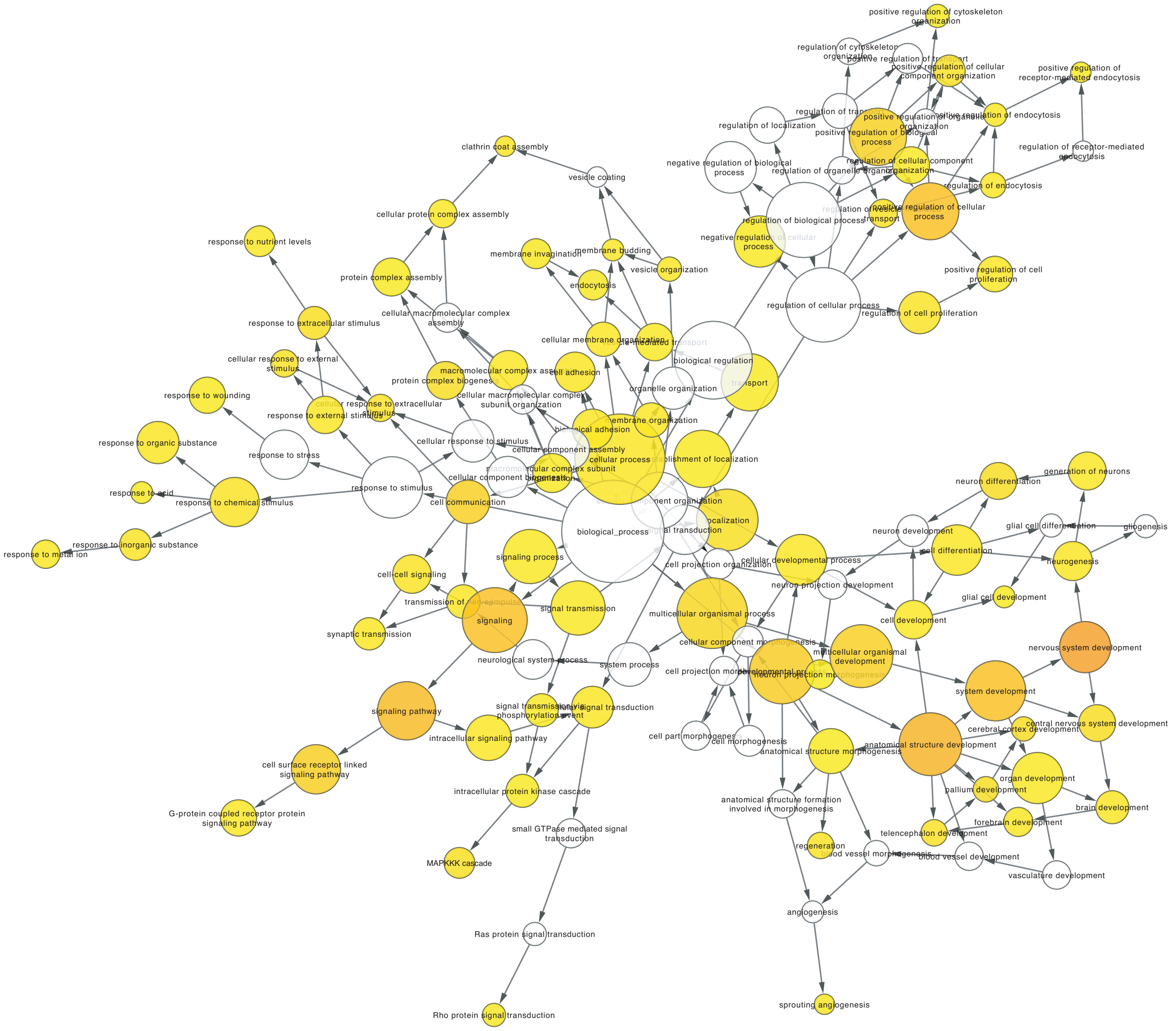


**Figure S6.** Gene Ontology (biological processes) network of differentially expressed genes by disease factor from BINGO in Cytoscape. The node size relates to number of genes, and the yellow nodes are statistically significant with a p-value $< 0.05$ and false discovery rate $< 0.05$.

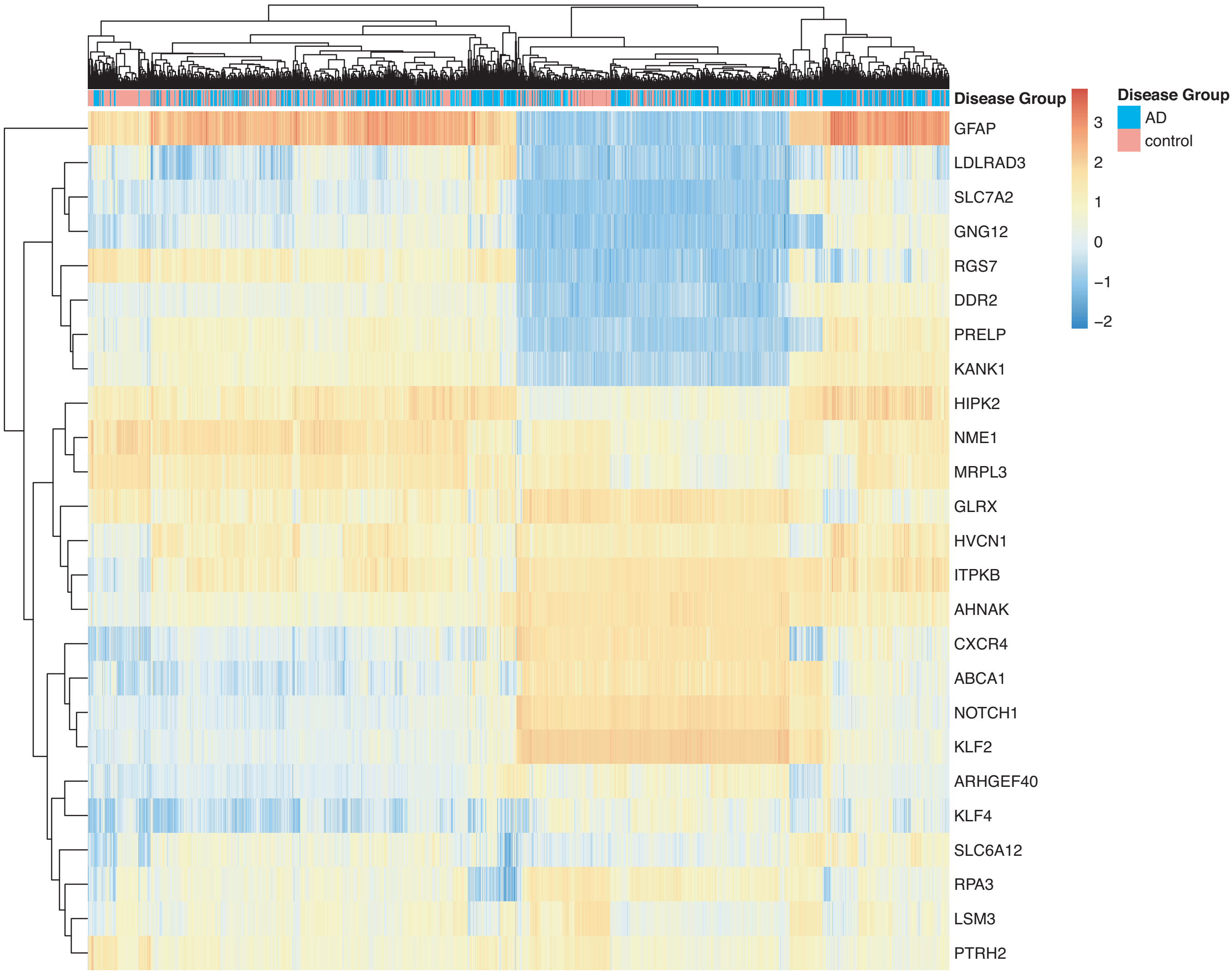


**Figure S7.** Heatmap with gene clustering of the top 25 differentially expressed disease (control-AD) gene list.

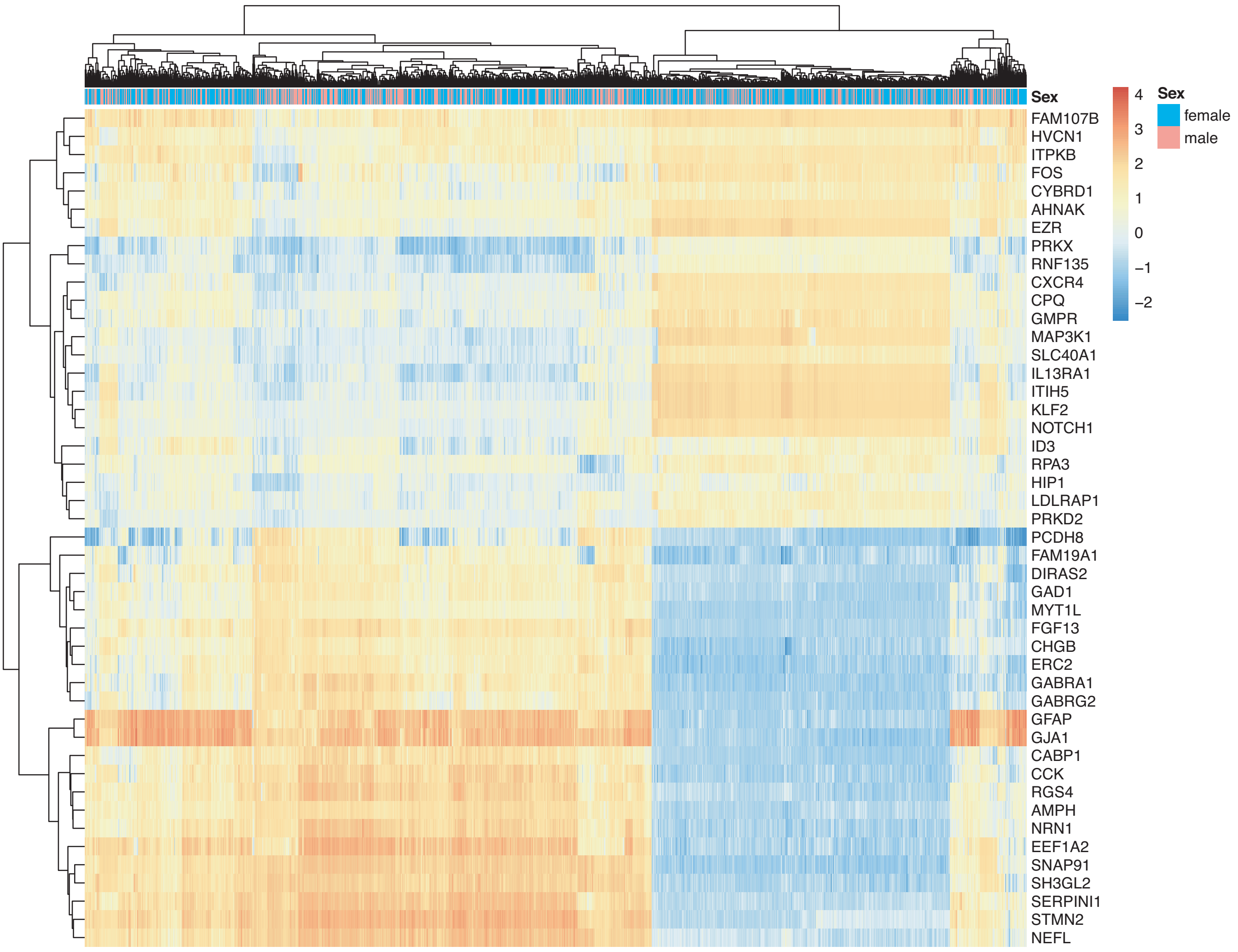


**Figure S8.** Heatmap with gene clustering to visualize gene expression in sex effect results on the differentially expressed disease (control-AD) gene list.

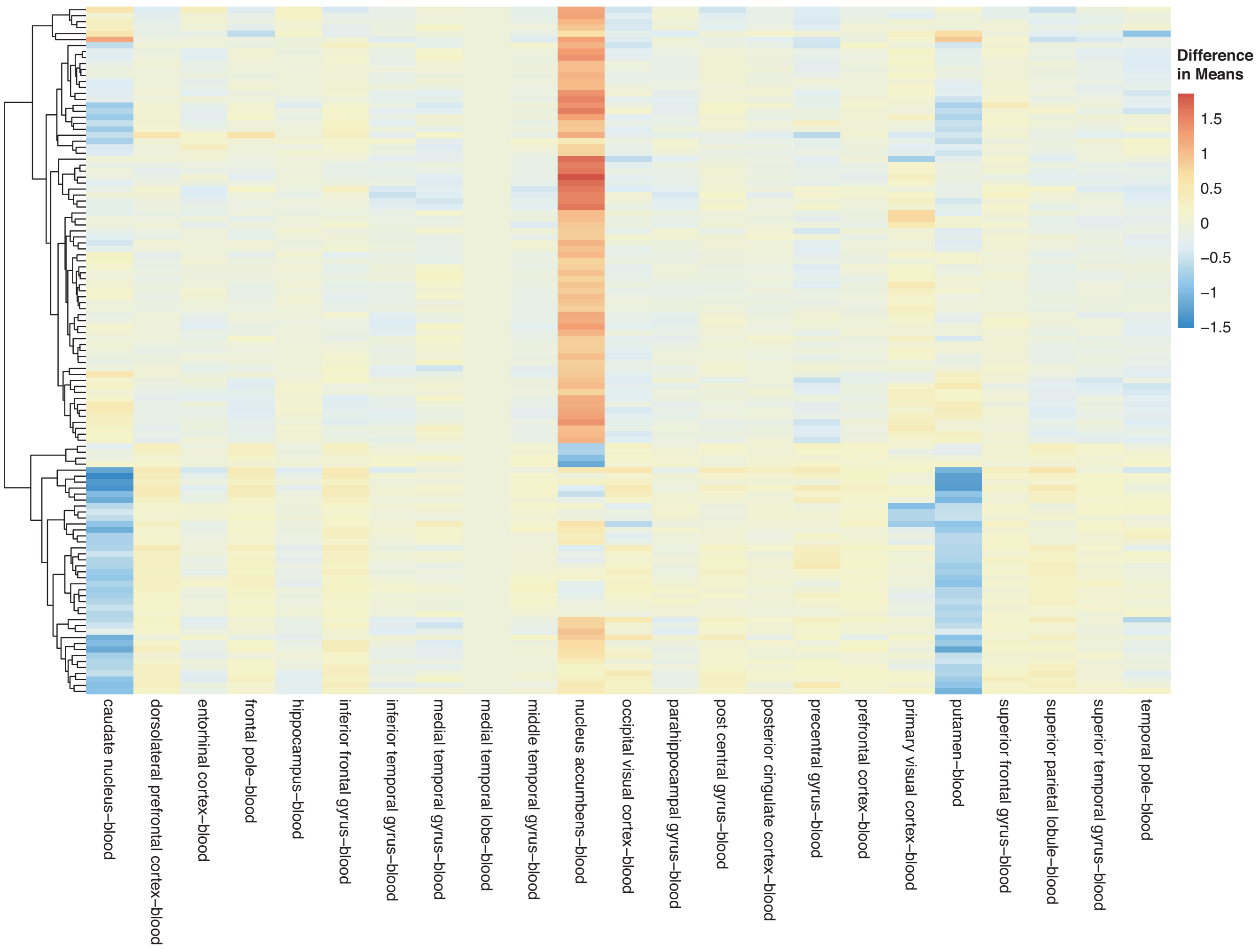


**Figure S9.** Heatmap with gene clustering to visualize tissue (blood as baseline) effect (differences in means between binary comparisons) on the differentially expressed disease (control-AD) gene list.
